# TMEM145 is a key component in stereociliary link structures of outer hair cells

**DOI:** 10.1101/2025.02.10.637577

**Authors:** Jae Won Roh, Kyung Seok Oh, Jiahn Lee, Yujin Choi, Soomin Kim, Ji Won Hong, Yelim Kim, Hogun Lew, Seung Hyun Jang, Hae-Sol Shin, Jiyeon Ohk, Hosung Jung, Kyoung Yul Seo, Jinwoong Bok, Chul Hoon Kim, Heon Yung Gee

## Abstract

Outer hair cells (OHCs) in the cochlea contain specialized stereociliary structures essential for auditory function. These include horizontal top connectors (HTCs), linking adjacent stereocilia and tectorial membrane-attachment crowns (TM-ACs), anchoring the tallest stereocilia to the tectorial membrane. The known molecular components of these structures, such as stereocilin, otogelin, otogelin-like, and tubby, lack transmembrane domains, suggesting the existence of anchoring proteins. This study identified TMEM145, a transmembrane protein with a Golgi dynamics (GOLD) domain, as a crucial OHC stereocilia component. TMEM145 was expressed in both OHCs and spiral ganglion neurons, with specific localization to TM-ACs and HTCs in OHCs. *Tmem145* knockout (KO) mice exhibited profound hearing impairment at three weeks of age, with complete loss of distortion product otoacoustic emissions, indicating OHC dysfunction. Immunostaining and scanning electron microscopy revealed the absence of TM-ACs and HTCs in *Tmem145* KO mice. In heterologous cell systems, TMEM145 interacted with stereocilin and tubby, facilitating their extracellular secretion. TMEM145 was undetectable in *stereocilin* KO and tubby mutant mice, indicating interdependence among these proteins. These findings establish TMEM145 as an essential membrane protein for the structural integrity of OHC stereocilia, providing insights into the molecular architecture of cochlear hair cells and their role in auditory function.

## INTRODUCTION

Hair cells in the cochlea are specialized sensory receptors essential for auditory signal transduction and convert mechanical sound vibrations into electrical signals^1, 2^. Among these, the hair bundles of outer hair cells (OHCs) play a crucial role in amplifying sound vibrations and refining the frequency resolution^2^. The structural integrity of hair bundles is crucial for detecting and responding to mechanical stimuli with high sensitivity. Therefore, a well-organized hair bundle with precisely arranged actin-rich stereocilia is necessary to maintain the efficiency of these auditory mechanisms^3^. Disruption of this organization because of genetic mutations can result in hearing impairment, a condition that affects millions of individuals worldwide^4, 5^.

OHC stereocilia are interconnected via means, such as tip links, horizontal top connectors (HTCs), and tectorial membrane-attachment crowns (TM-ACs), which maintain the well-organized structure and functional integrity of the sensory hair cells^6, 7^. Tip links connect the lateral wall of taller stereocilia to the tips of shorter ones and play a crucial role in opening mechanoelectrical transduction (MET) channels by transmitting the tension generated from stereocilia movement^8^. HTCs are zipper-like structures that interconnect adjacent stereocilia within and across rows, thereby contributing to the structural cohesion and stability of the stereocilia bundle. TM-ACs anchor the tallest row of stereocilia to the tectorial membrane (TM), ensuring proper mechanical coupling for auditory signal transmission. Various molecular components involved in the formation of these links have been identified, including PCDH15, CDH23, STRC, OTOG, OTOGL, and TUB, most of which are encoded by genes associated with hereditary deafness^9, 10, 11, 12, 13, 14^. Mutations in these genes have been shown to impair link integrity, leading to disorganization of the hair bundle structure, which results in hearing loss^8, 12, 15, 16, 17^.

Among the proteins involved in hair cell function, stereocilin (STRC), Otogelin (OTOG), otogelin-like (OTOGL), and tubby (TUB) are key contributors to the formation and maintenance of TM-ACs and HTCs. Mutations in *STRC*, *OTOG*, and *OTOGL* have been associated with autosomal recessive nonsyndromic hearing loss DFNB16, DFNB18B, and DFNB84, respectively,^12^ underscoring their importance in auditory physiology. However, none of these proteins possess a transmembrane domain, raising important questions about how they are secreted, transported, and anchored to the stereociliary membrane to maintain the integrity of stereociliary links. Identifying core proteins that possess transmembrane domains and interact with the OTOG(OTOGL)-STRC-TUB will be crucial for understanding the formation and maintenance of stereociliary links.

To identify transmembrane proteins involved in the auditory process, we used the International Mouse Phenotyping Consortium (IMPC) database^18^ (https://www.mousephenotype.org/) to search for candidate genes associated with deafness. This search led to the identification of the TMEM145 protein, which consists of a Golgi dynamics (GOLD) domain and a seven-transmembrane domain. A *Tmem145* knockout mouse model exhibited profound hearing impairment. Although the specific role of TMEM145 in auditory physiology remains unclear, these findings suggest that it plays a crucial role in the structural and functional integrity of cochlear hair cells.

The presence of a conserved GOLD domain in TMEM145 is of particular interest. The GOLD domain, first identified in the wntless (WLS) protein, is known for its role in facilitating protein trafficking and secretion. In WLS, the GOLD domain mediates the binding and transport of WNT ligands, which are crucial for cellular signaling and development^19, 20, 21^. Other GOLD domain–containing proteins, such as TMEDs/p24 and FYCO1, have also been implicated in vesicular transport and intercellular communication^22^. These observations highlight the versatility of the GOLD domain in mediating interactions between intracellular and extracellular environments.

Given the established role of the GOLD domain in other systems, we hypothesized that TMEM145 is a candidate anchor protein for the lateral links of stereocilia, utilizing its GOLD domain to interact with or regulate the localization of proteins essential for stereociliary links, such as STRC and TUB. In this study, we aimed to investigate the functional relationship among TMEM145, STRC, and TUB, with a focus on elucidating the potential role of TMEM145 in mediating these interactions. By integrating genetic, biochemical, and structural approaches, we provides evidence for the central role of TMEM145 in coordinating the assembly and precise localization of STRC and TUB within OHC stereocilia.

## RESULTS

### Predicted structure of TMEM145

We searched for deafness-associated candidate genes in the IMPC database^18^ to identify proteins that had not yet been implicated in auditory functions. The identification of these genes could potentially lead to the discovery of previously unreported deafness-associated genes in humans^23^. We concentrated on genes predicted to encode single or multiple transmembrane domains because we believed that these proteins could potentially act as novel receptors that had not been previously identified in auditory signal transduction. Among these candidates, *Tmem145* gained our attention as it was predicted to contain seven transmembrane domains^24^ (Supplementary Fig. 1a); moreover, its function had not been previously investigated.

First, we aimed to gain functional and structural insights by analyzing the predicted structure of TMEM145. According to predictions generated by AlphaFold3^25^, TMEM145 comprises an N-terminal domain, followed by seven transmembrane domains. Sequence analysis revealed that the first 29 amino acids at the N-terminus serve as a signal peptide (Fig. 1a, b)^26^. The DALI server^27^ was used to assess structural similarities between the TMEM145 domains and known domains within the human proteome predicted by AlphaFold2^28, 29^. Analysis of the N-terminal domain revealed a significant similarity to the Golgi dynamic (GOLD) domain found in TMED proteins and the SEC14-like protein family (Fig. 1b, Supplementary Fig. 1b, and Supplementary Table 1). The GOLD domain is well known for its roles in protein trafficking and secretion^19^. Further analysis of the seven transmembrane domains revealed significant similarities to class A or F G protein-coupled receptors (Fig. 1b, Supplementary Fig. 1c, and Supplementary Table 2). Comparison of the full-length structure of TMEM145 with other predicted structures revealed that the most significant matches were with seven transmembrane proteins containing GOLD domains—WLS, TMEM87A, TMEM87B, TMEM181, GPR107, GPR108, and GPR180 (Fig. 1b, Supplementary Fig. 1d, and Supplementary Table 3). Among these homologs, WLS is the most thoroughly characterized owing to its crucial function in the trafficking and secretion of WNT proteins^20,30, 31^. TMEM87A has recently been implicated in protein trafficking and ion channel activity^32, 33, 34, 35^. Other members of this family have also been reported to participate in trafficking pathways^36, 37, 38, 39^. Collectively, these findings indicate that TMEM145 may possess structural and functional similarities to proteins involved in protein trafficking and secretion.

**Fig. 1.**
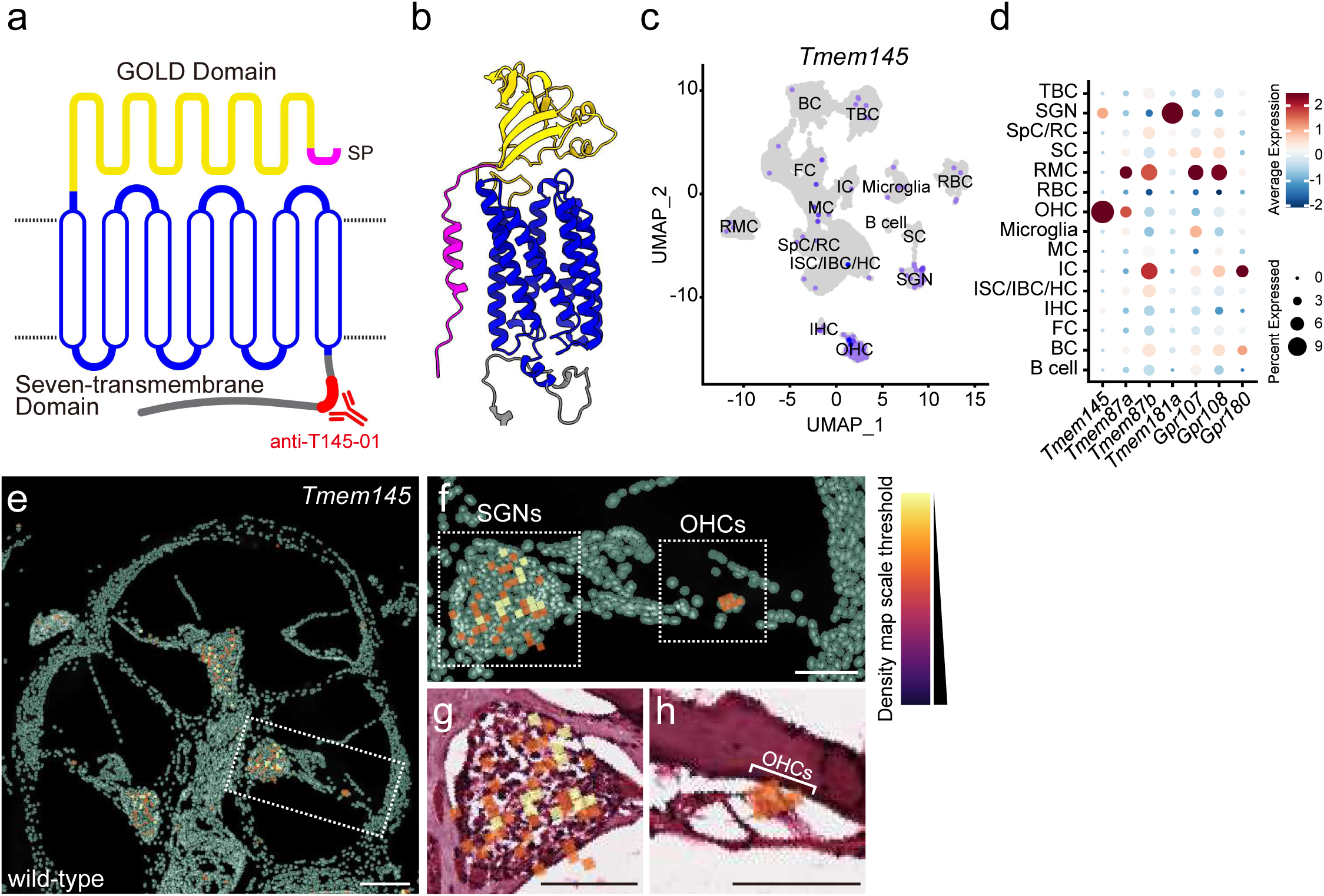
*Tmem145* is expressed in the outer hair cells of cochlea. **a.** Schematic of TMEM145, showing the N terminal GOLD domain (yellow), seven predicted transmembrane segments (blue), and the signal peptide (pink). The epitope recognized by the custom antibody (antiT14501) is indicated in red near the Cterminus. **b.** AlphaFold3^25^ predicted three dimensional structure of full length TMEM145, color coded to highlight the GOLD domain (yellow), seven transmembrane bundle (blue), signal peptide (magenta), and additional linker regions (gray). **c.** UMAP projection of single cell RNA sequencing data, illustrating *Tmem145* expression patterns in various cochlear cell types. Each dot represents an individual cell type and the intensity of the purple shading denotes *Tmem145* expression level. TBC, Tympanic border cell; SGN, Spiral ganglion neuron; SpC, Spindle cell; RC, Root cell; SC, Schwann cell; RMC, Reissner’s membrane; OHC, Outer hair cell; MC, Marginal cell; IC, Intermediate cell; ISC, Inner sulcus cell; IBC, Inner border cell; HC, Hensen cell; IHC, Inner hair cell; FC, Fibroblast; BC, Basal cell. **d.** Dot plot comparing the average expression levels of *Tmem145* and other related genes (*Tmem87a*, *Tmem87b*, *Tmem181a*, *Gpr107*, *Gpr108*, and *Gpr180*) across multiple cochlear cell populations. The color scale reflects mean expression, and the size of each dot indicates the proportion of cells expressing the gene of interest. **e.** Representative Xenium-based spatial transcriptomics density map of *Tmem145* expression type mouse cochlea cross section, visualized with a color scale from low (black) to high (yellow). Regions of interest are marked by dashed boxes. **f.** Enlarged view of the areas outlined in (**e**), focusing on spiral ganglion neurons (SGNs) and outer hair cells (OHCs). **g, h.** Higher magnification histological sections corresponding to the same areas shown in positive cells (orange) against hematoxylin/eosin (pink). Scale bars: (**e–h**) 200 μm.

### Expression of *Tmem145* in the organ of Corti

Subsequently, we examined the expression of *Tmem145* in mouse organ of Corti. Single-cell RNA sequencing (scRNA-seq) on postnatal day 28 (P28) revealed that *Tmem145* was mostly expressed in outer hair cells (OHCs); however, expression was also detected in spiral ganglion neurons (SGNs)^40^ (Fig. 1c, d, Supplementary Fig. 2). To confirm these findings, we conducted spatial transcriptomics, which indicated *Tmem145* expression in OHCs and a subset of SGNs, consistent with scRNA-seq results (Fig. 1e-h, Supplementary Fig. 3). SGNs are categorized as type I, which innervate inner hair cells (IHCs) and type II, which innervates OHCs^41^. Examination of SGNs categorized from the scRNA-seq data revealed that *Tmem145* exhibited higher expression in type I SGNs (Supplementary Fig. 4)^41^. Collectively, these results indicate that *Tmem145* is highly expressed in OHCs and a subset of SGNs, particularly type I SGNs, in adult mice.

### Auditory phenotypes of *Tmem145* knockout (KO) mice

Next, we investigated the role of *Tmem145* in auditory function. For this, we generated *Tmem145* KO mice (*Tmem145^-/-^*) by deleting exons 2–12 of *Tmem145*, located on chromosome 7 of the C57BL6/N strain, using CRISPR-Cas9 (Supplementary Fig. 5). The *Tmem145^−/−^* strain was viable and fertile, producing offspring in Mendelian ratios. We then assessed hearing phenotypes by measuring the auditory brainstem response (ABR). *Tmem145^−/−^* mice exhibited severe hearing impairment across all tested frequency ranges by P21, with hearing thresholds significantly increased by 4 weeks of age (Fig. 2a–c). Comparison of wave I amplitude, which represents the synchronized neural response of the auditory nerve to sound stimuli originating from the organ of Corti, revealed that *Tmem145^−/−^* mice had a markedly reduced amplitude relative to wild-type (WT) controls, indicating a defect in hair cell–mediated sound transmission (Fig. 2d).

**Fig. 2.**
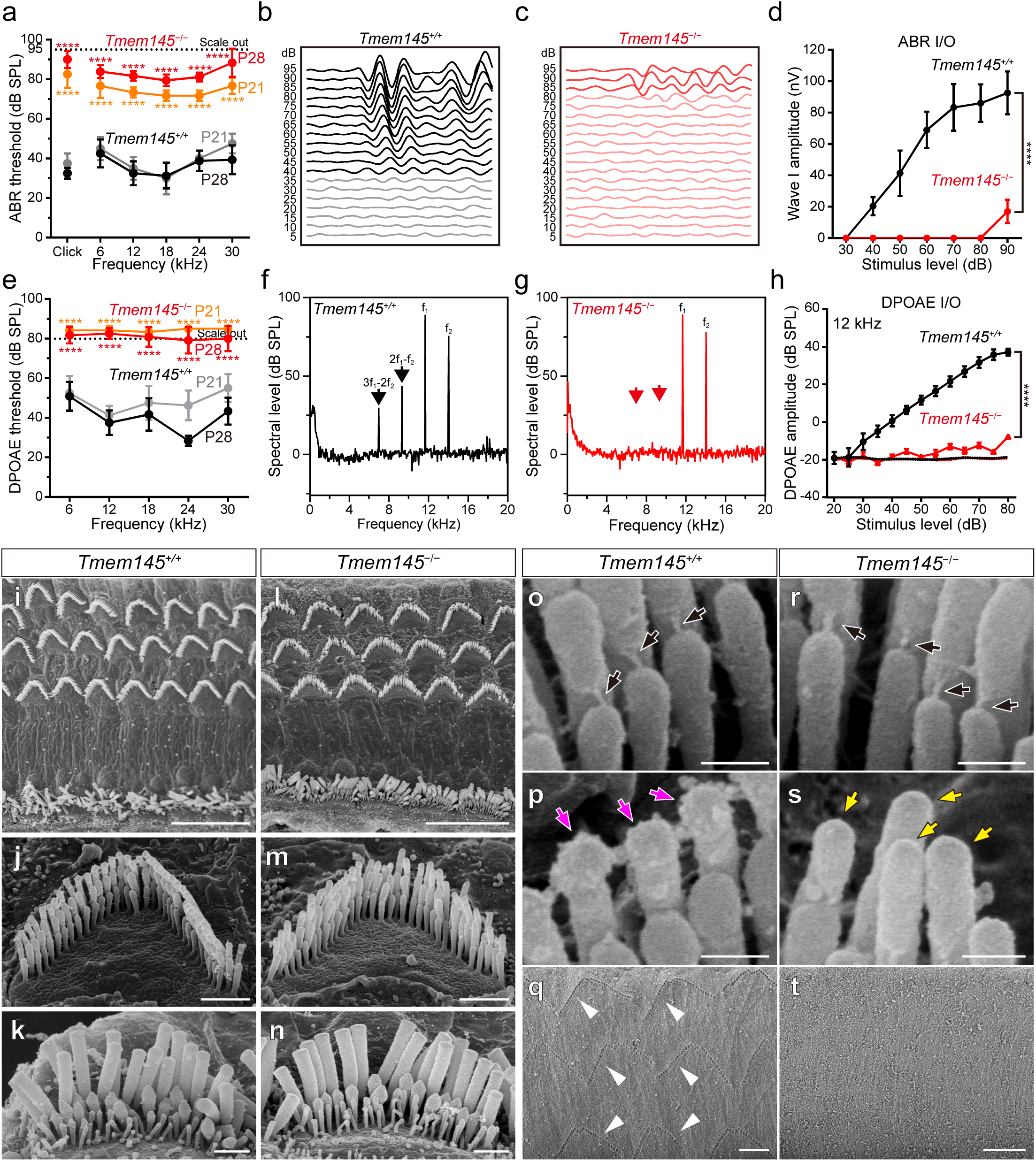
TMEM145 is essential for auditory function and the structural integrity of TM-ACs and HTCs in OHC stereocilia. **a.** Auditory brainstem response (ABR) thresholds at P21 and P28 in *Tmem145^+/+^* and *Tmem145^−/−^* mice. Red symbols for *Tmem145^−/−^* mice indicate significantly elevated thresholds, with some values reaching the upper limit (‘scale out’) of the measuring apparatus. n = 4 for *Tmem145^+/+^* P21; 8 for *Tmem145^+/+^* P28; 6 for *Tmem145^−/−^* P21; 9 for *Tmem145^−/−^* P28. **b**, **c.** Representative ABR waveforms at varying stimulus intensities in *Tmem145^+/+^* (**b**) and *Tmem145^−/−^* (**c**) mice. Each trace represents a 5-dB increase. **d.** ABR input–output (I/O) curves, showing the wave I amplitude as a function of stimulus level in *Tmem145^−/−^* and *Tmem145^+/+^* mice at P28. n = 8 for *Tmem145^+/+^*; 9 for *Tmem145^−/−^*. **e.** Distortion product otoacoustic emission (DPOAE) thresholds measured across multiple frequencies in *Tmem145^+/+^* and *Tmem145^−/−^* mice at P21 and P28. n = 4 for *Tmem145^+/+^* P21; 6 for *Tmem145^+/+^* P28; 6 for *Tmem145^−/−^* P21; 6 for *Tmem145^−/−^* P28. **f**, **g.** Frequency spectra of DPOAEs in *Tmem145^+/+^* (**f**) and *Tmem145^−/−^* (**g**) mice at a representative stimulus level. Black and red arrows indicate peaks at f_1_, f_2_, and their distortion products (2f_1_-f_2_ and 3f_1_-2f_2_) in *Tmem145^+/+^* and *Tmem145^−/−^* mice, respectively. **h** DPOAE I/O function at 12 kHz, demonstrating reduced responses at P28 in *Tmem145^−/−^* compared with *Tmem145^+/+^* mice. n = 6 for *Tmem145^+/+^*; 6 for *Tmem145^−/−^*. **i–n** Scanning electron microscopy (SEM) images of OHCs in *Tmem145^+/+^* (**i–k**) and *Tmem145^−/−^* (**l–n**) mice at P21. **o**, **p**, **r**, **s** Higher magnification SEM images of OHC stereocilia in *Tmem145* (**o**, **p**) and *Tmem145^−/−^* (**r**, **s**) at P21. Black arrows indicate preserved tip links. TM-ACs and HTCs are present in *Tmem145^+/+^* (**p**, magenta arrows), however, absent in *Tmem145^−/−^* (**s**, yellow arrows). **q**, **t** SEM images of imprints on the undersurface of TM in *Tmem145^+/+^* (**q**) and *Tmem145^−/−^* (**t**) mice at P21. White arrows indicate TM imprints in *Tmem145^+/+^*mice. Scale bars: (**i–n**) 20 µm; (**o–t**) 5 µm. Statistical analyses in (**a**) and (**e**) were conducted using two-sided unpaired t-tests by comparing at the same postnatal day. Statistical analyses in (**d**) and (**h**) were conducted using two-way analysis of variance followed by Dunnet’s post hoc test. ****p < 0.0001. Data are presented as mean ± standard error of mean.

To further investigate the underlying cause of hearing impairment, we performed distortion-product otoacoustic emission (DPOAE) measurements. DPOAEs are cochlear-generated sound emissions that reflect OHC electromotility, with abnormalities indicating impaired OHC-mediated sound amplification and cochlear dysfunction. *Tmem145^−/−^* mice displayed no detectable DPOAE signals across all frequency ranges at P21 (Fig. 2e–g). The analysis of DPOAE amplitude in relation to input stimulus revealed a significant reduction in *Tmem145^−/−^* mice (Fig. 2h). In contrast, heterozygous (Het) mice showed normal hearing phenotypes, indicating that *Tmem145* is not haploinsufficient in the auditory system (Supplementary Fig. 7a, b).

Next, we aimed to elucidate the mechanisms underlying hearing loss mediated by OHC defects. Three primary causes were considered: (1) loss of OHCs, (2) impaired mechanical coupling between OHCs and cochlear structures, such as the tectorial membranes, which could disrupt sound transmission, and (3) direct or indirect disruption of Prestin, the motor protein essential for OHC-driven cochlear amplification^42^. We performed scanning electron microscopy (SEM) on OHCs from P21 *Tmem145^+/+^* and *Tmem145^−/−^* mice to analyze these possibilities. No discernible loss of OHCs was observed in *Tmem145^−/−^* mice at low magnification (Fig. 2i, l). Slight loss of OHCs was observed only at P112 in *Tmem145^−/−^* mice, suggesting that OHC viability was not the major pathological outcome of TMEM145 loss (Supplementary Fig. 7). High-magnification SEM images showed disorganized OHC stereocilia bundles in *Tmem145^−/−^* mice (Fig. 2j, m), whereas IHC stereocilia remained structurally intact (Fig. 2k, n). Based on these findings, we further examined the ultrastructure of OHC stereocilia. Strikingly, while the tip link was preserved in *Tmem145^−/−^* mice (Fig. 2o, r), the TM-ACs and HTCs in OHCs were completely absent in *Tmem145^−/−^* mice (Fig. 2p, s). Furthermore, SEM analysis revealed the compete loss of the typical TM imprint in *Tmem145^−/−^* mice (Fig. 2q, t), phenocopying the *Strc* KO and tubby mice^17^. Voltage-clamp recordings of Prestin activity in OHCs showed no differences between WT and KO mice (Supplementary Fig. 8). The preservation of TM-AC is essential for the connection between tallest stereocilia and TM^17^; thus, the absence of TM-AC and HTC in *Tmem145^−/−^* mice strongly suggests that their disruption plays a major role in the observed hearing impairment.

### TMEM145 localization in the OHC stereocilia

Next, we examined the localization of TMEM145 in OHCs. To validate TMEM145 protein expression, we generated a TMEM145-specific antibody targeting the C-terminal region of the protein (Fig. 1a; anti-T145-01). The evaluation of this antibody in a heterologous system validated its specificity for both human and mouse TMEM145 (Supplementary Fig. 5). Immunostaining of organ of Corti samples from P21 *Tmem145^+/+^* and *Tmem145^−/−^* mice using anti-T145-01 revealed distinct signals in WT tissue, whereas negligible signals were seen with *Tmem145^−/−^* tissue (Fig. 3a, e). Consistent with the SEM findings, we observed substantial disruption in the arrangement of stereocilia in *Tmem145^−/−^* mice at P21 (Fig. 3e).

**Fig. 3.**
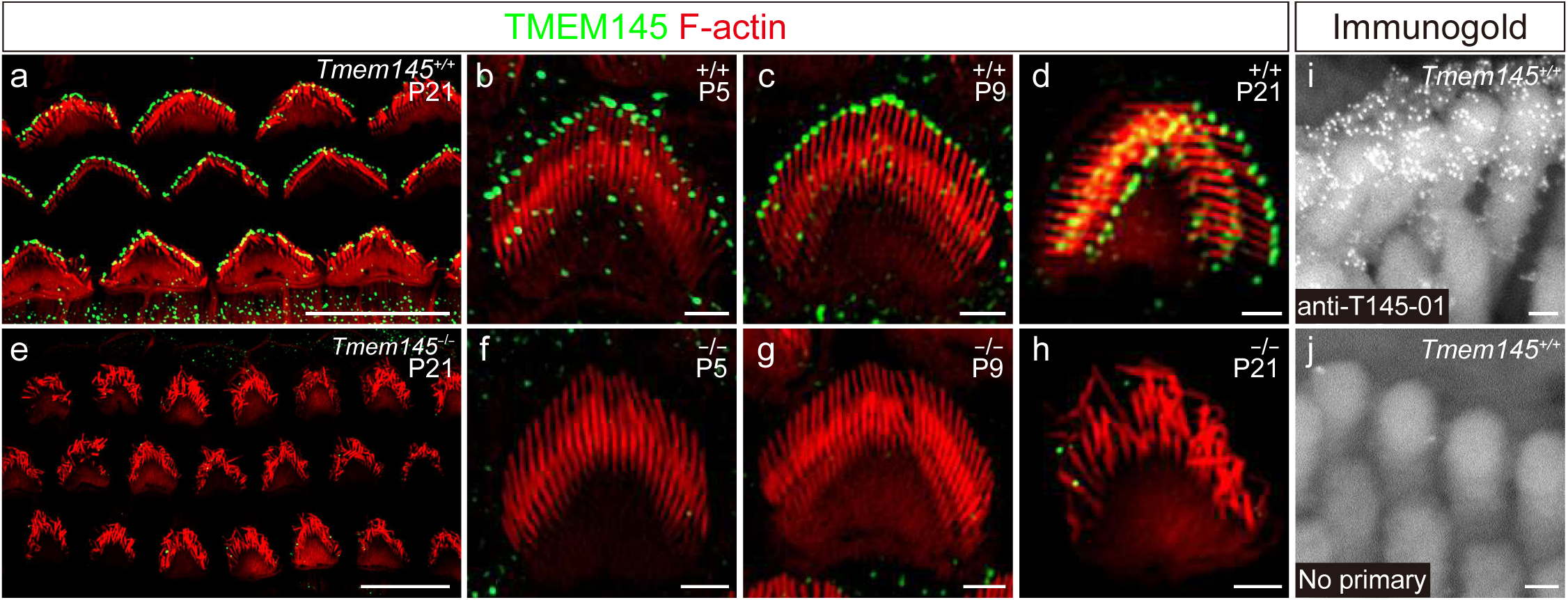
TMEM145 expression in OHC stereocilia and its association with TM-ACs and HTCs. **a–d.** Whole-mount immunofluorescence images of stereocilia from *Tmem145^+/+^* mice at P5, P9, and P21. Green signals represent TMEM145 (anti-T145-01), and red signals represent F-actin (phalloidin). (**a**) shows whole-mount immunostaining at P21, and (**b–d**) are high-magnification views of single outer hair cells at P5, P9, and P21, respectively. In the WT, TMEM145 is evenly distributed along the tips of stereocilia at all stages. **e–h.** Whole-mount immunofluorescence images of *Tmem145^−/−^* mice at the same time points. (**e**) displays whole-mount immunostaining at P21. (**f–h**) High-magnification images at P5, P9, and P21. In the mutant mice, TMEM145 signals are absent, and the stereocilia exhibit disorganized morphology and arrangement compared with controls. Scale bars: (**a**, **e**) 20 µm; (**b–d**, **f–h**) 5 µm.

A detailed analysis of whole-mount preparations from WT mice at a higher magnification revealed that TMEM145 is localized at the tips of OHC stereocilia, where at the TM-ACs and the HTCs are present (Fig. 3a, d). Sub-immunogold labeling using electron microscopy further confirmed its localization in these structures, providing strong evidence that TMEM145 is specifically enriched at TM-ACs and HTCs of OHC stereocilia (Fig. 3i, j).

To determine when stereocilia disorganization occurs during development, we subsequently examined earlier postnatal stages. In mice, OHC stereocilia undergo maturation until P14^6, 43^. Analysis of stereocilia morphology and TMEM145 localization at P5 and P9 revealed that TMEM145 was already present at the tips of tallest stereocilia starting from P5 onward (Fig. 3b, c). This expression pattern of TMEM145 aligns with that of STRC and TUB^17^. However, the stereocilia arrangement in *Tmem145*^−/−^ mice remained comparable to that of WT mice at both P5 and P9 (Fig. 3f, g). This suggests that while TMEM145 is present at the tips of tallest stereocilia from early postnatal stages, it is primarily required for preserving the stereocilia organization after they have fully matured.

### TMEM145 interacts with STRC and TUB, facilitating secretion

Next, we investigated the relationship between TMEM145 and other molecular components of TM-ACs and HTCs. Although STRC, OTOG, OTOGL, and TUB have been identified as essential components for TM-ACs and HTCs^15, 16, 17, 44^, none of these proteins possess a transmembrane domain. Given the membrane-bound nature, we hypothesized that TMEM145 functions as a linker and carrier, facilitating the localization and trafficking of these proteins to the OHC stereociliary membrane.

To test this hypothesis, we investigated whether TMEM145 physically interacts with other TM-AC components. Because of the low expression levels of TMEM145 and TM-AC proteins in Human Embryonic Kidney (HEK) 293T cells, we heterologously expressed C-terminal FLAG-tagged *TMEM145* in HEK293T cells, thereby confirming its robust expression (Supplementary Fig. 9). A cell surface biotinylation assay confirmed the presence of TMEM145 in the plasma membrane (Supplementary Fig. 9). Furthermore, immunostaining of overexpressing HeLa cells revealed that TMEM145 was localized to the endoplasmic reticulum (ER), Golgi apparatus, and plasma membrane (Supplementary Fig. 9).

We then introduced C-terminal MYC- or V5-tagged *STRC* and *TUB* under conditions in which each protein was co-expressed with or without *TMEM145*, followed by immunoprecipitation (IP) using FLAG beads. Co-immunoprecipitation (co-IP) of STRC and TUB with TMEM145 revealed physical interactions between TMEM145 and both proteins (Fig. 4a). Subsequent IP experiments using V5 beads to pull down TUB while observing the co-IP of TMEM145 and STRC demonstrated that TMEM145 co-immunoprecipitated with TUB, thus reaffirming a physical interaction (Fig. 4a). Notably, STRC coimmunoprecipitated with TUB only in the presence of TMEM145 (Fig. 4a). Considering that TUB is a cytosolic protein anchored to the membrane and that STRC (with a signal peptide) can localize to the ER lumen or extracellular space, these findings strongly suggest that TUB, TMEM145, and STRC form a physical association in the sequence TUB–TMEM145–STRC.

**Fig. 4.**
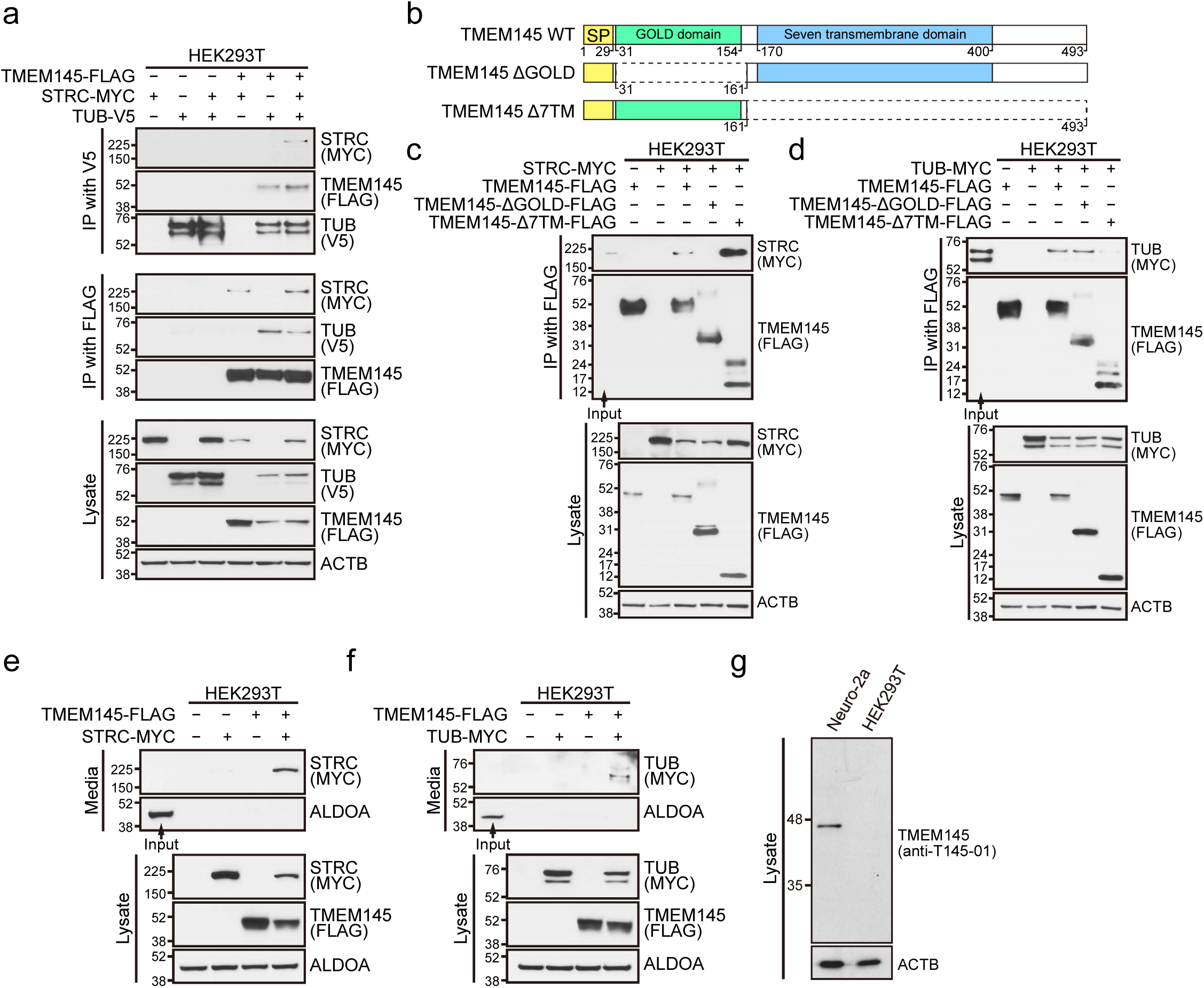
TMEM145 interacts with STRC and TUB and induces their secretion. **a.** HEK293T cells were transfected with *TMEM145*-FLAG, *STRC*-MYC, or *TUB*-V5, followed by immunoprecipitation (IP) using anti-V5 or anti-FLAG affinity matrices. The co-immunoprecipitated proteins (MYC, FLAG, V5) and total cell lysates were analyzed using immunoblotting. Under conditions where TMEM145-FLAG was co-expressed, STRC-MYC and TUB-V5 co-immunoprecipitated, indicating a physical interaction between TMEM145 and STRC/TUB. β-actin (ACTB) was used as a loading control. **b.** Schematic of TMEM145 wild-type (WT) and its deletion mutants (ΔGOLD, Δ7TM). SP indicates the signal peptide (amino acids 1–29). The GOLD domain spans amino acids 31– 154 and the seven-transmembrane domain is from amino acids 170–400. ΔGOLD lacks the GOLD domain, whereas Δ7TM lacks the seven-transmembrane region. **c.** HEK293T cells were co-transfected with STRC-MYC and either TMEM145-FLAG, TMEM145-ΔGOLD-FLAG, or TMEM145-Δ7TM-FLAG. Following IP with anti-FLAG, the presence of STRC-MYC in the immunoprecipitates was assessed using immunoblotting with anti-MYC. Deletion of the GOLD domain affects the interaction of TMEM145 with STRC, suggesting that those domains contribute to STRC binding. **d.** Similar co-transfections and immunoprecipitations were performed using TUB-MYC in place of STRC-MYC, with TMEM145-FLAG, TMEM145-ΔGOLD-FLAG, or TMEM145-Δ7TM-FLAG. Deletion of the 7TM domain impaired the interaction of TMEM145 with TUB. **e, f.** To examine the effect of TMEM145 on the secretion of STRC (**e**) and TUB (**f**). Culture supernatants (media) and cell lysates were analyzed using immunoblotting. Secreted STRC or TUB in the media was detected using immunoblotting, and ALDOA served as a loading control for both media and lysates. Both STRC and TUB were secreted only when TMEM145 was co-transfected. **g** Endogenous TMEM145 was detected in Neuro-2a and HEK293T cells using an anti-T145-01 antibody, with ACTB (β-actin) serving as a loading control.

Next, we attempted to identify which region of TMEM145 mediates its interactions with STRC and TUB. Using full-length TMEM145 as a template, we generated two GOLD and Δ7TM (Fig. 4b). ΔGOLD lacks the GOLD domain, however, retains the signal peptide and seven-transmembrane domain, whereas Δ transmembrane domain, however, contains the GOLD domain. The co-expression of these constructs with C-terminal MYC-tagged *STRC* followed by FLAG IP showed that STRC 7TM, however, not with ΔGOLD, indicating that the GOLD domain of TMEM145 is essential for binding to STRC (Fig. 4c). TUB exhibited the contrary behavior, binding only to ΔGOLD and not to Δ7TM, suggesting that the seven-transmembrane domain of TMEM145 mediates TUB binding (Fig. 4d). Collectively, these data indicated that TUB interacts with the transmembrane region of TMEM145 on the cytosolic side, whereas STRC binds to the GOLD domain on the extracellular or luminal side of TMEM145.

Finally, we investigated whether TMEM145 facilitates the secretion of STRC and TUB, similar to the role played by WLS in the trafficking and secretion of WNT^30, 31^. In OHCs, STRC is thought to be secreted to form the extracellular structures of TM-AC and HTC^16^. However, to our knowledge, STRC secretion has never been conclusively proven by showing a substance that regulates its secretion. Co-expression of TMEM145 and STRC in HEK293T cells resulted in the secretion of STRC into the culture medium, whereas STRC remained undetectable extracellularly in the absence of TMEM145 (Fig. 4e). The same phenomenon was observed for TUB (Fig. 4f).

The overexpression of TUB in Neuro-2a cells has been shown previously to lead to its secretion^45, 46^, whereas it is not in HEK 293T cells in the present study. This discrepancy prompted us to investigate whether endogenous TMEM145 was responsible. Using the anti-T145-01 antibody to detect TMEM145 levels, we discovered that TMEM145 was present in Neuro-2a cells, however, absent in HEK293T cells, suggesting that cell type-specific secretion of TUB could be mediated by endogenous TMEM145 (Fig. 4g).

Overall, these *in vitro* findings confirmed the physical interactions among TMEM145, STRC, and TUB, as well as the role of TMEM145 in promoting the secretion of STRC and TUB, similar to the mechanism by which WLS facilitates WNT secretion.

### Structural interactions among proteins that comprise the ultrastructure of OHC stereocilia

Considering the previously demonstrated interactions among TMEM145, STRC, and TUB in a heterologous system, we examined how these elements affect each other in OHC stereocilia. Utilizing validated antibodies from our earlier study^17^, we examined the localization of STRC and TUB in *Tmem145^+/+^* and *Tmem145^−/−^* mice. At P5, 9, and 21, both STRC (Fig. 5a-h) and TUB (Fig. 5i-p) were undetectable in TM-ACs and HTCs of *Tmem145^-/-^* mice.

**Fig. 5.**
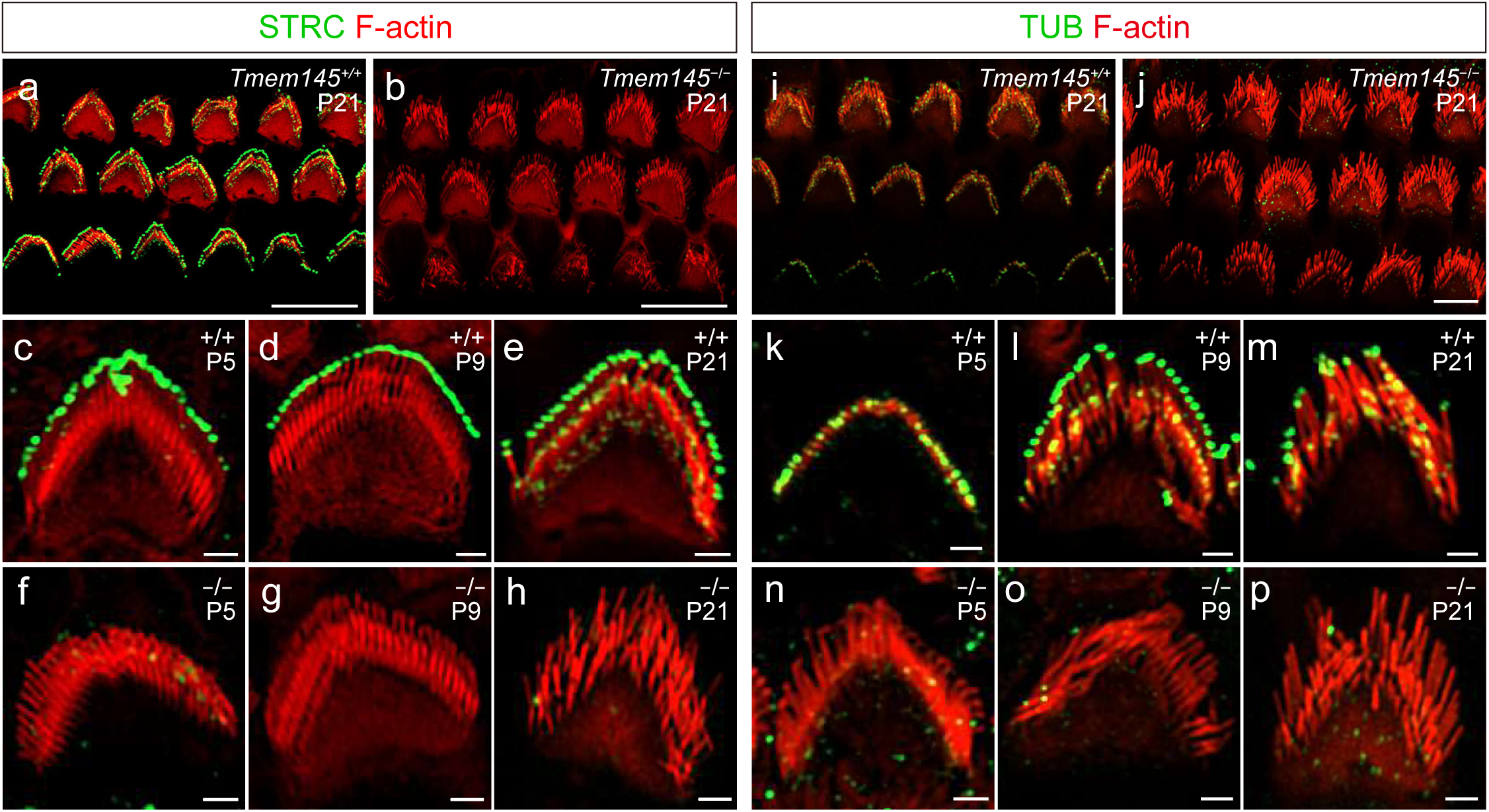
Mislocalized STRC and TUB in *Tmem145^−/−^* OHC stereocilia. **a, b.** Whole-mount immunofluorescence images of the organ of Corti at P21. Green signals represent STRC, and red signals represent F-actin (phalloidin). **c–h.** Developmental immunofluorescence images of STRC localization in OHC stereocilia in *Tmem145^+/+^* and *Tmem145^−/−^* mice at P5, P9, and P21. Compared with *Tmem145^+/+^* mice, STRC signals were absent in the OHC stereocilia of *Tmem145^−/−^* mice. **i**, **j.** Whole-mount immunofluorescence images of the organ of Corti at P21. Green signals represent TUB, and red signals represent F-actin (phalloidin). **k–p.** Developmental immunofluorescence images of TUB localization in OHC stereocilia in *Tmem145^+/+^* and *Tmem145^−/−^* mice at P5, P9, and P21. Compared with *Tmem145^+/+^* mice, TUB signals were absent in the OHC stereocilia of *Tmem145^−/−^* mice.

We then assessed TMEM145 localization in mouse strains with genetic disruption of other proteins essential for the ultrastructure of OHC stereocilia (Fig. 6a). TMEM145 was absent in the stereocilia of *Strc^−/−^* and *Otog^−/−^* mice (Fig. 6b, c). To study the role of *Tub*, we used two different genetic backgrounds. In *tub/tub;Map1a^B6^* mice, in which neither TM-AC nor HTC was observed at P21^17^, TMEM145 was similarly lacking (Fig. 6d). In *tub/tub;Map1a^AKR^* mice, where TM-AC was present at P21, however, HTC was not^17^, TMEM145 was found exclusively at TM-AC (Fig. 6e).

**Fig. 6.**
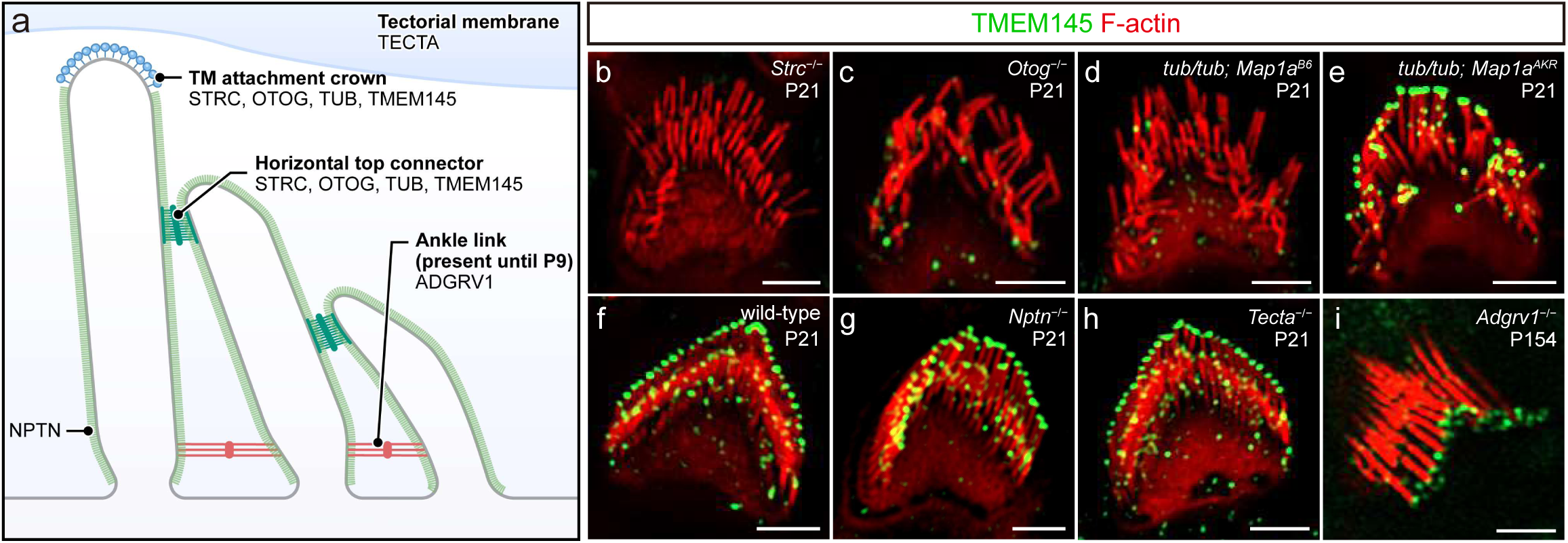
Localization of TMEM145 is only affected in TM-AC and HTC components. **a.** Schematic showing the overall structure of OHC stereocilia. The localization of the proteins used in each mouse line is indicated. **b–i.** Whole-mount images of OHC stereocilia from P21 mice immunolabeled for TMEM145 (green) and F-actin (red). Panels depict (**b**) *Strc*^−/−^, (**c**) *Otog*^−/−^, (**d**) *tub/tub; Map1a*^B6^, (**e**) *tub/tub; Map1a*^AKR^, (**f**) wild-type, (**g**) *Nptn*^−/−^, (**f**) *Tecta*^−/−^, and (**i**) *Adgrv1*^−/−^ genotypes. Each panel shows stereocilia morphology and the localization of TMEM145. All images are shown at similar magnification, with scale bars indicating 5 µm.

To determine whether TMEM145 mislocalization was exclusively because of the absence of the TM-AC or HTC components, we examined TMEM145 localization in multiple mouse strains deficient in proteins unrelated to these structures (Fig. 6a). These included NPTN, which is present throughout the stereocilia and modulates calcium pumps^47^; TECTA, which is specifically localized to the TM and maintains its integrity^48, 49^; and ADGRV1, which establishes ankle links in immature stereocilia^50, 51^. In all three KO lines— *Nptn^−/−^*, *Tecta^−/−^*, and *Adgrv1^−/−^*—TMEM145 localization was unaffected (Fig. 6f–i).

Collectively, these results demonstrate that in *Tmem145^−/−^* mice, the normal localization of key TM-AC and HTC components was disrupted, whereas genetic disruption in these components also caused TMEM145 mislocalization. This interdependence underscores the interrelated functions of TMEM145, STRC, TUB, OTOG, OTOGL, and other TM-AC/HTC proteins in the formation and preservation of the OHC stereociliary structures.

## DISCUSSION

In this study, we investigated TMEM145, an uncharacterized protein containing a seven-transmembrane protein and GOLD domain. *Tmem145^−/−^* mice exhibited severe hearing loss at three weeks of age owing to compromised TM-ACs and HTCs in OHCs. Biochemical experiments revealed that TMEM145 physically interacted with STRC and TUB, while also mediating their secretion (Fig. 7a). *In vivo* analyses further showed that the absence of TMEM145 impaired the localization of STRC and TUB, and the disruption of these proteins similarly affected TMEM145 localization (Fig. 5 and Fig. 6). These findings suggest the potential of TMEM145 as a molecular scaffold for assembling TM-ACs and HTCs and stabilizing their components during or after secretion.

**Fig. 7.**
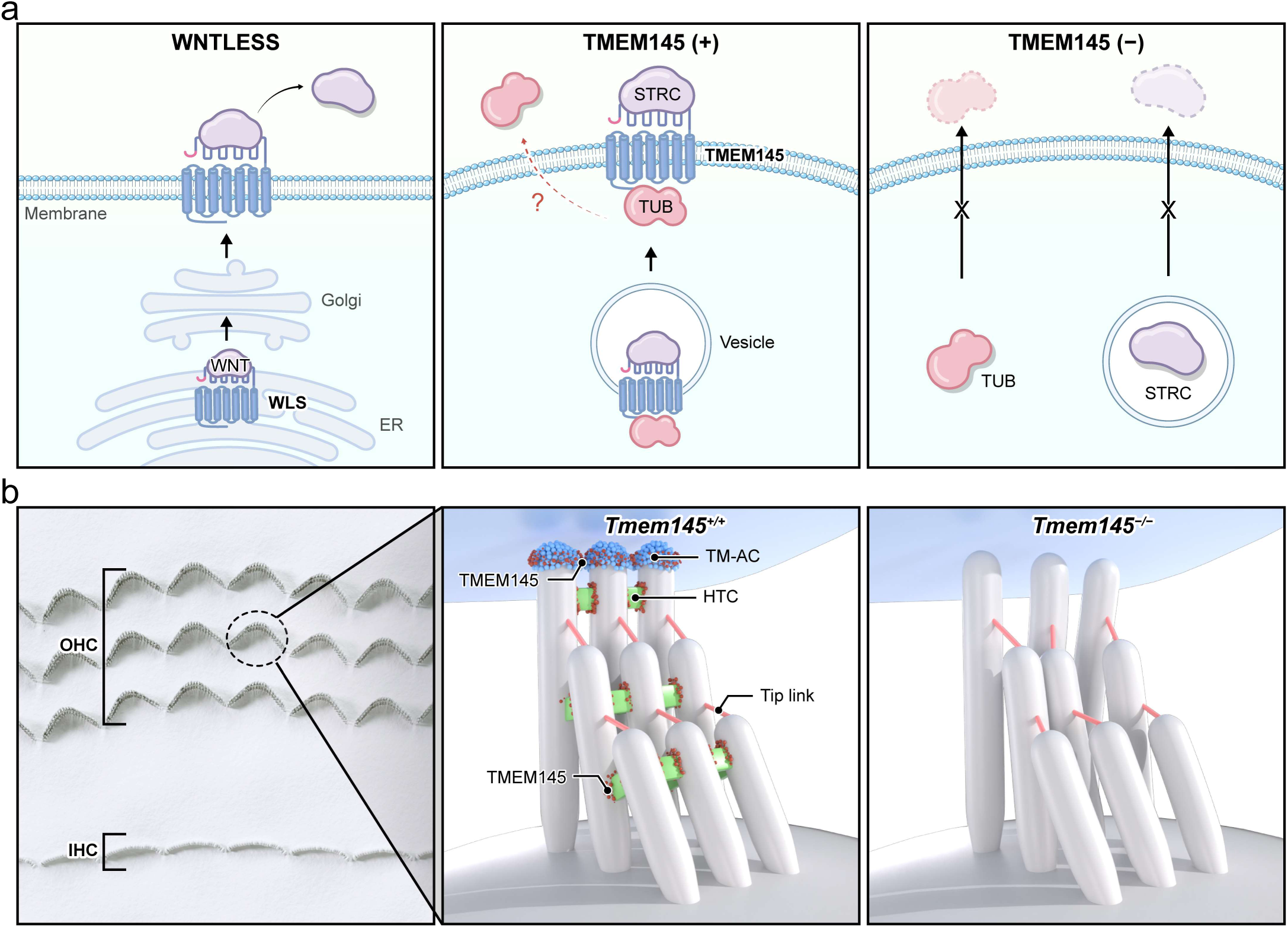
Graphical illustration of TMEM145 function. **a.** WNT is trafficked from the endoplasmic reticulum and Golgi to the cell surface with the help of WLS (left panel), and TMEM145 coordinates the proper trafficking of stereociliary proteins, such as STRC and TUB (middle panel). When TMEM145 is absent, these proteins fail to reach the designated location. **b** A top view of organ of Corti with three rows of OHC and one row of IHC (left panel). The 3D illustrations compare *Tmem145^+/+^* (middle panel) and *Tmem145^−/−^* (right panel) OHC stereocilia, showing that TMEM145 deficiency cause the loss of TM-AC and HTC with tip link intact and disrupts stereociliary alignment.

Previous studies have suggested that STRC is probably secreted at the tips of the tallest stereocilia based on the observation that it was detected on the imprints of the lower side of the TM, even when the TM was mechanically detached from the stereocilia^16, 17^. The identification of TMEM145 as a regulator of STRC secretion is an important step in confirming that STRC is a secreted protein. However, the maintenance of STRC at the tip of stereocilia even after secretion raises a new question about its mechanism. The first possibility is that TM anchors STRC to the stereocilia tip and prevents detachment. However, *Tecta^−/−^* mice showed the presence of STRC even in the absence of attachment between stereocilia and TM, dismissing the possibility^16^ (Supplementary Fig. 10). Throughout this study, the relationship between STRC and TMEM145 was noted similar to that between WNTs and WLS in the WNT secretion pathway (Fig. 7a). Assuming that the two pairs of molecules share similar regulatory principles, STRC may sit on TMEM145, a transmembrane protein, and dissociate from TMEM145 only when the concentrations of certain molecules are sufficient to displace STRC from TMEM145, just as secreted FZD-related protein and WNT inhibitory factor 1 release WNTs from WLS in the plasma membrane^31^. The presence of the corresponding extracellular molecules in the endolymph remains unknown. Revealing the identities and changes in various physiological and pathological conditions will further our understanding of the functional stability of secreted STRC in maintaining hearing.

We found that TUB localization to the tip of the stereocilia also requires STRC, suggesting that STRC and TUB are mutually essential for the proper localization of both proteins to the tip of the stereocilia (Supplementary Fig. 11). In addition, OTOG and OTOGL are absent at the tip of stereocilia in *Strc^−/−^* mice^15^. The results of this study have further revealed the complexity of these interactions by showing that STRC, TUB, and OTOG all require TMEM145 for their localization to the stereocilia, and vice versa (Fig. 5 and Fig. 6). These seemingly reciprocal interactions between the proteins constituting a stereociliary ultrastructure make it difficult to determine which protein among them plays a more primary and central role. In our previous study^17^, we observed the transient localization of STRC to the tip of stereocilia at P5 in tubby mice; however, STRC was not present in the stereocilia at P9 or later. This was an unexpected pattern. The disappearance of both proteins in the stereocilia of *Tmem145^-/-^* mice at P5 suggests that TMEM145 is the central protein that regulates other secreted proteins, even in immature OHCs. This early organization driven by TMEM145 would allow the establishment of more sophisticated regulatory mechanisms in mature OHCs, where each component is strictly interdependent for hair cell functions.

The interactions of these proteins and their arrangement in quaternary structures remain unclear. The information we obtained in this study indicates that TUB and STRC do not directly interact with each other, however, that TMEM145 bridges them by interacting with each protein through different domains on TMEM145. The nature of the interactions of OTOG with other proteins remains unknown. However, immunostaining experiments for TUB, STRC, and OTOG provided some clues. To stain these proteins, the TM should be mechanically detached to expose the proteins at the tips of the stereocilia. Although STRC or TUB usually stick to the lower side of the TM in some amounts, in most cases, we can successfully visualize the proteins at the tips of the stereocilia in an even distribution.

Conversely, OTOG and OTOGL are not consistently detected at the tips of the OHC hair bundles, with varied immunostaining intensities between the experiments. *Tecta^−/−^* mice, where TM is inherently detached from the cochlear epithelium, showed consistent immunostaining of OTOG and OTOGL proteins^15^, indicating weak adherence of OTOG to the tips of stereocilia during TM detachment. Furthermore, *Otog^−/−^* and *Otogl^−/−^* mice showed markedly weaker STRC immunostaining signals at the tips than WT mice; however, a significant signal intensity remained^15^. This is in contrast to experiment where OTOG and OTOGL were not present in *Strc^−/−^* mice^15^. These data suggest that OTOG and OTOGL may be weakly, however, directly associated with TMEM145 or indirectly associated with TMEM145 via TUB or STRC, maintaining TM-ACs.

Our study also presents future opportunities to elucidate the regulation and function of TUB, which remain poorly understood. TUB has been reported to undergo unconventional secretion in Neuro-2a, a neuronal cell line^45^. However, its mechanism of action remains unknown. The well-known molecular features of TUB are to have nuclear localization signal at its N-terminus and a phosphatidylinositol-4,5-bisphosphate (PI(4,5)P_2_)-binding domain at its C-terminal region^52^. However, binding of TUB to PI(4,5)P_2_ has only a partial effect on its secretion from Neuro-2a^45^. Confirmation of TMEM145 expression in Neuro-2a cells (Fig. 4g) suggested that TMEM145 may be a major regulator of TUB secretion in Neuro-2a and other cell types. In terms of function, TUB is a MerTK ligand that facilitates the phagocytosis of retinal pigmental epithelial cells, macrophages, and microglia^46, 53^. MerTK is a member of the TAM receptor protein tyrosine kinase family (Tyro3, Axl, and MerTK) and its expression on macrophages plays pivotal roles in innate immunity. Although macrophages are not present in the endolymph, they are abundant in the endolymphatic sac, which is connected to the endolymph^54^. The bone marrow–derived resident macrophages play important roles in immune defense and prevent inflammation of the sensory epithelium^54^. Secreted TUB in the endolymph may reach the endolymphatic sac and activate MerTK in macrophages. Although we could not find an immunophenotype in tubby or *Tmem145^−/−^* mice, the roles of TUB in inner ear immunity may be revealed in tubby or *Tmem145^−/−^* mice in pathological conditions, such as infection or acoustic trauma.

Detection of TMEM145 expression in spiral ganglion neurons (SGNs), particularly in type I SGNs, suggests that its role in auditory physiology extends beyond stereocilia maintenance in OHCs. Given its structural features—an N-terminal GOLD domain linked to protein trafficking and a seven-transmembrane domain reminiscent of class A G protein-coupled receptors—TMEM145 may be integral to synaptic function in SGNs. It can potentially regulate the trafficking or localization of synaptic proteins, influence synaptic vesicle cycling, and participate in intracellular signaling pathways essential for neural signal transmission. This dual expression profile raises the possibility that TMEM145 contributes to both the structural integrity of hair cell stereocilia and proper functioning of neuronal synapses, thereby ensuring efficient auditory processing and communication along the auditory pathway.

Overall, this study provides compelling evidence for the central role of TMEM145 in coordinating the assembly and precise localization of STRC and TUB within OHC stereocilia, offering a new perspective on the molecular mechanisms underlying auditory transduction. Future research on the detailed dynamics of TMEM145-mediated protein trafficking and stereociliary organization will significantly deepen our understanding of the cellular processes that support hearing physiology.

## METHODS

### Mice

*Tmem145* knockout mice were generated by Cyagen (Suzhou, China). The *Tmem145* gene (NCBI Reference Sequence: NM_183311.3; Ensembl: ENSMUSG00000043843) contains 15 exons, with an ATG start codon in exon 1 and a TGA stop codon in exon 15 (Transcript Tmem145-201: ENSMUST00000108409). Exons 2–12 were selected as target regions. Cas9 protein and two gRNAs (gRNA-A1: GCCAGGCAACCTCCACTTGG-TGG and gRNA-A2: GGCTTTCACGGAGCTAGGCA-AGG) were co-injected into fertilized mouse eggs. The injected embryos were then transferred into pseudopregnant females, and the resulting F0 offspring were screened for mutations. Genotyping was performed using PCR and the knockout allele was confirmed using Sanger sequencing.

### TMEM145 antibody production

A synthetic peptide corresponding to the sequence GVPGPGGSQSADK located at the C-terminus of human and mouse TMEM145 was commercially synthesized and subsequently conjugated to a carrier protein, keyhole limpet hemocyanin (KLH), to enhance its immunogenicity. This peptide–carrier conjugate was prepared following established cross-linking protocols.

New Zealand white rabbits were immunized with the peptide conjugates according to a standard immunization regimen. An initial primary injection was administered followed by booster injections at regular intervals. However, additional boosting did not result in a substantial increase in the immune response. Consequently, no further boosting was performed, and final serum collection was performed based on the immune response obtained from the initial immunization series.

Blood was collected from the rabbits by standard venipuncture and the serum was separated by centrifugation. Sera obtained from individual animals were pooled to generate a uniform final serum sample. The pooled serum was subjected to Protein A affinity chromatography to purify the rabbit polyclonal antibodies. The final antibody preparation was formulated with Proclin 300 and stored at −80 °C for subsequent use. The final antibody product was designated as *anti-T145-01*. The purified antibody was used at a 1:200 dilution for whole-mount immunostaining and a 1:1000 dilution for immunoblotting.

### Auditory brainstem response (ABR)

ABR thresholds were measured in a soundproof chamber using Tucker-Davis Technologies (TDT) RZ6 digital signal processing hardware and BioSigRZ software (Alachua, FL, USA). Subdermal needles (electrodes) were positioned at the vertex and ventrolateral to the right and left ears of the anesthetized mice. A calibrated click stimulus (10 burst stimuli (5 ms duration) were produced at 4, 6, 8, 10, 12, 18, 24, 30, and 42 kHz using SigGenRZ software and an RZ6 digital signal processor and were delivered to the ear canal by a multifield 1 (MF1) magnetic speaker (TDT). The stimulus intensity was increased from 10 to 95 dB SPL in 5-dB increments. The ABR signals were fed into a low-impedance Medusa Biological Amplifier System (RA4LI, TDT), which delivered the signal to the RZ6 digital signal-processing hardware. The recorded signals were filtered using a 0.5–1 kHz bandpass filter and the ABR waveforms in response to 512 tone bursts were averaged. ABR thresholds for each frequency were determined using BioSigRZ software. Peak amplitudes (mV) and peak latencies (ms) were calculated from the waveform signals of the click-evoked ABRs as input/output (I/O) functions with increasing stimulus levels (20–90 dB SPL).

### Distortion product otoacoustic emissions (DPOAE)

DPOAE were measured using a TDT microphone–speaker system. Primary stimulus tones were produced using an RZ6 digital signal processor with SigGenRZ software and delivered using a custom probe with an ER 10B+ microphone (Etymotic, Elk Grove Village, IL, USA) and MF1 speakers positioned in the ear canal. The primary tones were set at a frequency ratio (f2/f1) of 1.2 with target frequencies of 6, 8, 10, 12, 16, 18, 24, and 30 kHz. The f2 intensity levels were the same as the f1 intensity levels (L1 = L2). The sounds resulting from the primary tones were received by the ER 10B+ microphone and recorded using an RZ6 digital signal processor. The DPOAE I/O functions were determined at specific frequencies (6 and 30 kHz) with a frequency ratio (f2/f1) of 1.2 and equal intensity levels (L1=L2). The intensity level of the primary tones was increased from 20 to 80dB SPL in 5-dB increments. Fast Fourier transform (FFT) was performed at each primary tone for the DP grams and at each intensity for the I/O functions using BioSigRZ software to determine the average spectra of the two primaries, the 2f1–f2 distortion products, and the noise floors.

### Immunofluorescence staining and whole-mount assay

The cochleae were isolated from wild-type, *Tmem145^−/−^*, *Strc^−/−^*, *Otog^−/−^*, *tub/tub;Map1a^B6^*, and *tub/tub;Map1a^AKR^* mice at P5, P7, and P21. Temporal bones were fixed with 4% paraformaldehyde overnight at 4 °C and decalcified in 250 mM EDTA for at least 2 d. The cochlea was dissected into pieces from the decalcified tissue for whole-mount immunofluorescence. Tissues were permeabilized with 0.3% SDS, blocked with 5% goat serum for 2 h, and then incubated at 4 °C overnight with the following primary antibodies: rabbit anti-T145-01, rabbit anti-Stereocilin, and mouse anti-Tubby. To label IHCs and OHCs, Alexa Fluor 594 phalloidin (1:400; A12381; Invitrogen) was used to stain F-actin at RT for 1 h. After three rinses with 1× PBS, the specimens were incubated for 1 h with the secondary antibodies goat anti-rabbit Alexa Fluor 488 (1:1000 dilution; A32731; Invitrogen) and goat anti-mouse Alexa Fluor 488 (1:1000 dilution; A32723; Invitrogen). The specimens were mounted in Faramount Aqueous Mounting Medium (S3025; Dako, Glostrup, Denmark) and imaged with a confocal microscope (LSM980; Zeiss, Jena, Germany) at 63x, and images were processed using ZEN software.

### Immunohistochemistry

The isolated cochleae were fixed with 4% paraformaldehyde overnight at 4 °C and decalcified in 250 mM EDTA for at least 2 d. The cochleae were transferred to a series of gradient ethanol solutions, immersed in dimethylbenzene, and embedded in paraffin for sectioning. For immunostaining, sections were deparaffinized in xylene and rehydrated in an ethanol gradient. The cells were then incubated in blocking buffer containing 5% goat serum for 1 h at room temperature. The samples were incubated overnight at 4 °C with primary antibodies diluted in blocking buffer. After washing with PBS, the samples were incubated with secondary antibodies and 4′,6-diamidino-2-phenylindole (DAPI) for 30 min at room temperature, washed, and covered with mounting medium and cover slips. Images were captured using a confocal microscope (LSM980; Zeiss).

### Scanning electron microscopy (SEM)

Inner ears were isolated in 1× PBS and initially fixed with 4% paraformaldehyde (PFA) for 40 min, followed by pre-fixation in a solution of 2.5% glutaraldehyde and 2% PFA in 0.1 M sodium cacodylate buffer containing 2mM calcium chloride for 1 h on ice. The cochlear epithelium was then micro-dissected in 1× PBS and further fixed overnight at 4 °C in a fixative comprising 2.5% glutaraldehyde, 2% PFA in 0.1 sodium a odylate buffer supplemented with 3.5% sucrose and 2 mM calcium chloride. Following fixation, the samples were washed three times with 0.1 M sodium cacodylate buffer (containing 2 mM calcium chloride) for 20 min each on a shaker maintained on ice. Post-fixation was carried out using an osmium tetroxide/thiocarbohydrazide (OTO) protocol, which included incubation in 1% osmium tetroxide (OsO) for 1 h on ice with shaking, treatment with 1% thiocarbohydrazide (TCH) for 10 min at room temperature, and a subsequent 30 min incubation in 1% OsO with shaking. The specimens were then dehydrated using a graded ethanol series (20%, 40%, 60%, 70%, and 80% for 7 min each at room temperature, followed by 90% ethanol overnight, 95% ethanol for 7 min, and 100% ethanol for 20 min) and dried using a critical point dryer (Leica EM CPD300, Wetzlar, Germany). Finally, the dried samples were coated with a 20-nm thick layer of platinum and imaged using a Schottky field-emission scanning electron microscope (JSM-7610F; JEOL, Tokyo, Japan).

### Sub-immunogold labeling for electron microscopy

Cochlear samples were isolated from the inner ears of mice and fixed in 4% PFA in Hank’s Balanced Salt Solution (HBSS) supplemented with 2 mM CaCl and 0.5 mM MgCl (pH 7.4) at room temperature for 40 min. The dissection buffer was HBSS.

For simultaneous blocking and permeabilization, the samples were incubated in 5% goat serum and 0.1% BSA in HBSS containing 0.3% saponin and 0.3% SDS) at room temperature for 1 h, and incubated with primary antibodies diluted 1:200 in the blocking and permeabilization solution overnight at 4 °C. The samples were rinsed once and washed three times for 15 min each with HBSS and then incubated with colloidal gold–conjugated secondary antibodies (12 or 18 nm, Jackson Immuno Research) diluted 1:50 in the blocking and permeabilization solution overnight at 4 °C. After incubation with the secondary antibody, the samples were rinsed once and washed three times for 15 min each with HBSS.

To prepare for fixation, the samples were rinsed twice with 0.1 M sodium cacodylate buffer (pH 7.2) and fixed in a solution of 10% glutaraldehyde and 4% PFA in 0.1 M sodium cacodylate buffer at 4 °C for at least 24 h without agitation. Postfixation of the samples was performed following the OTOTO protocol. The samples were washed three times for 10 min with 0.1 M sodium cacodylate buffer and fixed with 1% osmium tetroxide (OsO) in 0.1 M sodium cacodylate buffer for 1 h at room temperature without agitation. The samples were washed four times for 5 min with 0.1 M sodium cacodylate buffer and then three times with distilled water. The samples were incubated with 1% thiocarbohydrazide for 20 min at room temperature without agitation, followed by four washes with water for 5 min each and three washes with 0.1 M sodium cacodylate buffer. This cycle was repeated to complete the OTOTO process.

Following the OTOTO protocol, samples were dehydrated in a graded ethanol series (15%, 30%, 50%, 75%, 90%, 95%, and 100%) for 5 min each on ice. The dehydrated samples were dried using a CPD300 critical point dryer (Leica Microsystems, Wetzlar, Germany) and coated with a 15-nm layer of carbon. Samples were imaged using a field-emission scanning electron microscope (JSM-IT800(); JEOL, Tokyo, Japan).

### Spatial transcriptomics

The Xenium workflow, utilizing experimental chemistry alongside a prototype instrument and consumables, started by sectioning 5-μm formalin-fixed paraffin-embedded tissue samples onto a Xenium slide. The sections were deparaffinized and permeabilized to expose the mRNA. The mRNA targets were detected using 313 probes, as described previously, and two negative controls: (1) probe controls to evaluate non-specific binding and (2) genomic DNA (gDNA) controls to confirm RNA-specific signals. Probe hybridization was carried out at 50 °C overnight at a concentration of 10 nM. Stringent washing followed to eliminate unbound probes, after which the probes were ligated at 37 °C for 2 h, during which an RCA primer was also annealed. Circularized probes were enzymatically amplified, first at 4 °C for 1 h and then at 37 °C for 2 h, producing multiple copies of gene-specific barcodes for each RNA binding event, enhancing the signal-to-noise ratio. Background fluorescence was chemically quenched to address the autofluorescence caused by lipofuscins, elastin, collagen, red blood cells, and formalin fixation. Finally, the tissue sections were mounted onto an imaging cassette and loaded onto a Xenium Analyzer.

### Single cell RNA sequencing analysis

Single-cell RNA-sequencing count matrices for 4-week-old mouse cochleae were obtained from the Gene Expression Omnibus (accession number GSE202920)^40^. All analyses were performed in R (v4.4.2) using Seurat (v5.1.0)^55, 56^. Feature barcode matrices from each sample were imported using Read10X and converted into Seurat objects. The two Seurat objects were merged into a single object to obtain a unified dataset.

For each cell, the percentage of reads mapping to mitochondrial genes was calculated using the PercentageFeatureSet with the pattern “^MT-”. Low-quality cells, such as those with few detected features or high mitochondrial gene content, were removed according to data-dependent thresholds. These thresholds included having between 200 and 3,000 detected features and fewer than 10% of the reads mapping to mitochondrial genes. Next, we used FindVariableFeatures (method = “vst”) to detect the 2,000 most highly variable genes across cells. Identifying highly variable genes helps focus on the genes that capture the greatest biological variability. To account for cell-cycle effects, we applied Seurat’s built-in cell-cycle scoring approach. Gene sets corresponding to the S phase (s_genes) and G2/M phase (g2m_genes) were used for CellCycle Scoring. Each cell was assigned scores for the S and G2/M phases and an overall phase annotation.

We employed the SCTransform workflow for normalization, which regularizes and stabilizes the variance in the single-cell data. We regressed out mitochondrial gene percentage (percent.mt) and cell cycle phase scores (Phase, S.Score, and G2M.Score) to mitigate their confounding effects on downstream analyses. Principal Component Analysis (PCA) was performed on the SCTransformed data using RunPCA, focusing on the variable features identified above. We typically selected the first 20 principal components (PCs) for downstream clustering, guided by inspection of an elbow plot or other metrics. Clustering was performed using a graph-based approach. First, a cell-to-cell similarity graph was constructed using FindNeighbors (dims = 1:20). We then applied Louvain or Leiden clustering using FindClusters (resolution = 1). We performed dimensional reduction via Uniform Manifold Approximation and Projection (UMAP) using RunUMAP (dims = 1:20). Clusters were visualized using DimPlot, and cells were colored according to their cluster identities. We identified cluster-specific marker genes using FindAllMarkers, applying a log-fold change threshold (e.g., 0.25) and focusing on positively differentially expressed genes (only.pos = TRUE). We annotated the clusters by examining known cell-type marker genes and the top differentially expressed genes. Based on these marker genes, we renamed the clusters using RenameIdents to reflect the known cochlear cell types. The final cell-type assignments were stored in the Seurat object metadata for downstream analysis.

To analyze single-cell RNA-sequencing data of SGNs, raw gene-by-cell count matrices for GSE114759^41^ were downloaded in a tab-delimited (*. txt) format, each containing gene symbols as row names and cell barcodes as column headers, and merged into a single matrix after retaining only genes common to all files. Low-quality cells (fewer than 100 detected genes) and genes detected in 10 or fewer cells were excluded. Cells expressing hemoglobin-related genes (Hba-a1, Hba-a2, Hbb-bh1, Hbb-bs, and Hbb-bt) were also excluded if they comprised over 5% of the total UMI count, thereby mitigating red blood cell contamination. The remaining data were log-normalized using Seurat’s NormalizeData function (scale factor 10,000) and 2,000 highly variable genes were identified using FindVariableFeatures. The expression matrix was then scaled (ScaleData), allowing for the optional regression of covariates, such as mitochondrial or cell cycle effects. PCA was conducted on the variable features and an elbow plot guided the selection of 20 PCs. These PCs informed graph-based clustering (FindNeighbors, FindClusters, resolution = 0.5) and two-dimensional visualization with UMAP (RunUMAP, dims = 1:20), which were used to inspect the cluster structure and annotate cell types based on known marker genes.

### Cell culture, plasmids, and transfection

HEK293T cells (CRL-3216; American Type Culture Collection, Manassas, VA, USA), HeLa cells (CCL-2; American Type Culture Collection), and Neuro-2a (CCL-131; American Type Culture Collection) were cultured in high-glucose Dulbecco’s modified Eagle’s medium (Thermo Fisher Scientific, USA) supplemented with 10% fetal bovine serum (Thermo Fisher Scientific) and 1% penicillin/streptomycin (Thermo Fisher Scientific). The cells were cultured in a humidified incubator at 37 °C with 20% O_2_ and 5% CO_2_, and were cultured every 2–3 d in 90-mm culture dishes.

Human *TMEM145* (NM_173633.3) and mouse *Tmem145* (NM_183311.3) coding regions were purchased from OriGene (Rockville, MD, USA). Each coding region was amplified using PCR to introduce a C-terminal FLAG tag (DYKDDDK) immediately upstream of the stop codon, and the fragments were subcloned into the pEGFP N1 mammalian expression vector (Clontech, USA) using the NheI and NotI restriction sites via In Fusion Cloning (Takara Bio, Japan). During this process, the original EGFP coding sequence was removed, leaving only the TMEM145 sequence under control of the CMV promoter.

The mouse *Strc* (NM_080459.2) and rat *Tub* (NM_013077.2) coding sequences were provided by Dr. Chul Hoon Kim (Yonsei University College of Medicine). Each coding region was amplified using PCR to introduce a C-terminal MYC tag (EQKLISEEDL) immediately prior to the stop codon. The resulting products were cloned into the pcDNA3.1(+) expression vector (Thermo Fisher Scientific) via In HindIII and NotI for mStrc and KpnI and XhoI for rTub. For some experiments, a tagged rTub (GKPIPNPLLGLDST) was used. The coding sequences of all plasmids were verified using Sanger sequencing (Macrogen, Seoul, South Korea).

HEK293T cells were seeded in six-well plates 1 d prior to transfection. Transient transfection was performed using Lipofectamine 2000 (Thermo Fisher Scientific) according to the manufacturer’s instructions. For single-protein expression, 1000 ng of either human or mouse TMEM145 plasmid was used. For co-expression experiments, the ratio and amount of each plasmid were optimized as follows: 200 ng TMEM145, 300 ng of TUB, and 3000 ng STRC plasmids. Molecular biology experiments were performed at 24–72 h post-transfection.

### OHC electrophysiology

OHC electrophysiology was performed as previously described with slight modifications^23^. The organ of Corti was acutely isolated from mice at P21. The animals were anesthetized using isoflurane (Sigma Aldrich) and euthanized by decapitation. The bone surrounding the apical turn of the cochlea was carefully removed and the apical turn of the organ of Corti was dissected using fine forceps and separated from the lateral cochlear wall, stria vascularis, modiolus, and tectorial membranes. All dissections were carried out in a standard extracellular solution containing 144 mM NaCl, 5.8 mM KCl, 10 mM HEPES, 5.6 mM glucose, 0.7 mM NaH PO, 1.3 mM CaCl, 0.9 mM MgCl, and 10 mM sorbitol, adjusted to pH 7.4 with NaOH. The isolated cochlea was immobilized on a 12-mm-diameter coverslip using a stainless-steel pin and Sylgard and mounted on an upright microscope (ECLIPSE FN1; Nikon, Japan). The extracellular bath solution used during recordings was identical to the dissection solution, while the pipette solution for recording electromotility contained 140 mM CsCl, 10 mM HEPES, 1 mM EGTA, 3 mM Mg ATP, and 1 mM MgCl, adjusted to pH 7.2 with CsOH.

Whole-cell voltage-clamp recordings were performed at 22–25 °C on OHCs using standard configurations. Microglass pipettes (World Precision Instruments, Sarasota, FL, USA) with a resistance of 2–5 MΩ were fabricated using a PP830 singlestage glass microelectrode puller (Narishige, Japan), and the liquid junction potential was corrected with an offset circuit prior to each recording. Currents were recorded using an Axopatch 700 B amplifier (Molecular Devices, San Jose, CA, USA) coupled with a Digidata 1550A (Molecular Devices) interface, digitized at 100 kHz, and low-pass filtered at 10 kHz using pClamp software 10.7 (Molecular Devices). The whole cell configuration was confirmed by ensuring that the series resistance remained below 10 MΩ, with no series resistance compensation applied.

In the voltage-clamp configuration, recordings were obtained using a sine wave stimulus (10 mV amplitude at 1 kHz) superimposed on a ramp pulse protocol ranging from 150 mV to 100 mV over a duration of 250 ms, with a holding potential of 70 mV. For each cell, at least two recordings were performed to ensure measurement stability, and the nonlinear capacitance (NLC) calculation was performed as described previously^23, 57, 58^. The capacitance was fitted to the derivative of the Boltzmann equation

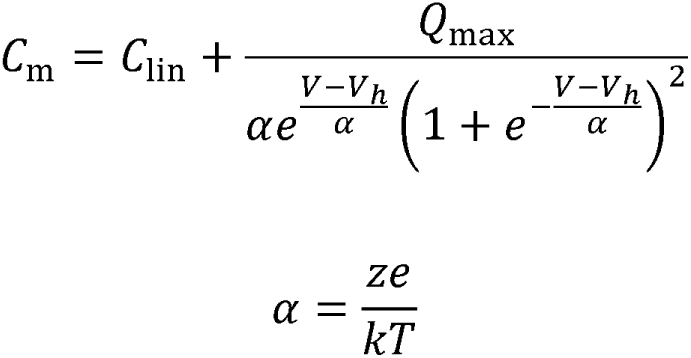

where *V_h_* is the maximal activation voltage, *Q_max_* is the maximal charge transfer between the plasma membrane, _α_ is the slope factor for the voltage-dependent charge transfer, *z* is the charge valence, *e* is the electron charge, *k* is Boltzmann’s constant, and *T* is absolute temperature. An in-house Python script was used for the NLC calculations and analysis^23^.

### Immunoprecipitation

Immunoprecipitation and co-immunoprecipitation experiments were performed as described previously^23^. Cells were processed and lysed under optimized conditions for the detection of surface and interacting proteins using a lysis buffer containing 150 mM NaCl, 50 mM Tris-Cl, 1 mM EDTA, 1% Triton X-100, and a Protease Inhibitor Cocktail (cOmplete Protease Inhibitor Cocktail tablets; Roche, Basel, Switzerland). The cell lysate was sonicated for 20 s and centrifuged at 20,000 × *g* for 15 min to remove cellular debris. Protein concentrations across samples were equalized using the Bradford assay (Protein Assay Kit II, Bio-Rad, Hercules, CA, USA). From each sample, a portion was used for immunoprecipitation, whereas the remainder was mixed with 2× Novex Tris-glycine SDS sample buffer (Sigma-Aldrich) and used as a lysate or input control. The lysate was then subjected to immunoprecipitation by incubation for overnight at 4 °C with EZview Red Anti-FLAG M2 Affinity Gel (Sigma-Aldrich), Pierce Anti-c-Myc Magnetic Bead (Thermo Fisher Scientific), or V5-Trap Magnetic Agarose (ChromoTek, Planegg, Germany) on an end-over-end rotator.

Following immunoprecipitation, the affinity matrices were washed five times with lysis buffer at a volume at least 20-fold greater than the matrix volume for each wash to remove nonspecifically bound proteins. For sample elution, 2× sample buffer (Sigma-Aldrich) was mixed with lysis buffer at a 1:1 ratio to achieve a final 1× concentration, and the mixture was incubated with the affinity matrix at 37 °C for 60 min. After elution, samples were collected in a new tube and subjected to immunoblotting.

### Surface biotinylation

Surface biotinylation was performed using EZ-Link Sulfo-NHS-SS-Biotin (Thermo Fisher Scientific) and streptavidin (Thermo Fisher Scientific) as previously described^59, 60^. The biotin solution was prepared by dissolving 0.5 mg EZ-Link Sulfo-NHS-SS-Biotin in 1 mL of 1× PBS. Cells were incubated in the dark at 4 °C for 30 min to allow for the biotin reaction. Subsequently, any remaining biotin was quenched by incubating the cells in a blocking solution containing 0.5% bovine serum albumin (prepared by dissolving 0.5 g BSA in 50 mL of 0.5× PBS) in the dark at 4 °C for 10 min. Following biotinylation, the biotinylated proteins were isolated using the immunoprecipitation procedure described above, with a Streptavidin agarose resin used to capture the proteins, followed by sample elution and immunoblotting.

### Immunoblotting

Lysate or input samples containing 40 μg of protein were loaded, while immunoprecipitation or biotinylation samples corresponding to the precipitated material from 400μg of lysate were loaded onto a 4–12% gradient SDS-PAGE gel (KomaBiotech, Seoul, Korea). Electrophoresis was performed at 100 V for 100 min using running buffer composed of 100mM Tris base, 100mM Tricine, and 0.1% SDS, adjusted to pH7.3. Proteins were then transferred onto an Amersham™ Protran® Supported nitrocellulose membrane (Cytiva, Marlborough, MA, USA) at 250 mA for 100 min using a transfer buffer containing 25 mM Tris, 192 mM Glycine, pH 7.3, and 20% methanol, with the transfer performed at 4 °C. The membrane was then blocked at room temperature for 1 h in 5% skim milk and incubated overnight at 4 °C with mild agitation, and the desired primary antibody diluted in a solution containing 5% skim milk and 0.02% sodium azide. The next day, the membrane was washed thrice for 10 min each at room temperature with 1× TBST composed of 20 mM Tris base, 150 mM NaCl, 0.1% Tween-20, adjusted to pH 7.4, incubated with the appropriate secondary antibody diluted in 1× TBST for 1 h at room temperature, and washed thrice for 10 min each with 1× TBST. Finally, immunoreactive bands were detected by developing the blot with ECL solution (Invitrogen) and visualized on a blue film (Agfa, Mortsel, Belgium).

For immunoblotting of secreted proteins from the culture media^61^, the medium was replaced with serum-free medium 24 h after transfection. The cells were then incubated for an additional 48 h before the culture medium was harvested. The harvested medium was concentrated using an Amicon Ultra-4 filter (Millipore, Burlington, MA, USA). Finally, the concentrated media were mixed with 5× sample buffer and equal volumes of each sample were loaded for subsequent analyses.

The following antibodies were used. Primary: anti-MYC (1:1000 dilution; 2276; Cell Signaling Technologies, Danvers, MA, USA), anti-FLAG (1:1000 dilution; 14793; Cell Signaling Technologies), anti-V5 (1:1000 dilution; R960-25; Invitrogen), anti-ACTB (1:1000 dilution; A284; Abbkine, Atlanta, Georgia, USA), and anti-ALDOA (1:1000 dilution; sc-390733, Santa Cruz Biotechnology, Santa Cruz, TX, USA). Secondary: goat anti-rabbit IgG polyclonal antibody (HRP conjugate) (1:1000 dilution; ADI-SAB-300-J, Enzo Life Sciences, Farmingdale, NY, USA) and goat anti-mouse IgG F(ab’)2 polyclonal antibody (HRP conjugate) (1:1000 dilution; ADI-SAB-100-J; Enzo Life Sciences).

### Immunocytochemistry

Transfected HeLa cells were cultured on 18-mm round coverslips. The cells were washed twice with 1× PBS, fixed with 4% PFA for 10 min, washed twice with 1× PBS, and permeabilized with 0.15% Triton X-100 in 1× PBS for 10 min. The cells were washed twice with PBS and incubated with blocking buffer consisting of 5% donkey serum for 1 h at room temperature. After blocking, the cells were incubated with the appropriate primary antibodies for 1 h at room temperature and washed thrice with 1× PBS. Subsequently, the cells were stained with secondary antibodies conjugated to a fluorescent marker for 1 h at room temperature and washed thrice with 1× PBS^62^. The samples were mounted onto glass slides using a mounting medium (Dako) and images were captured using a confocal microscope (LSM 700; Zeiss) equipped with a 40× objective lens.

### Structural modeling

The full-length structure of human TMEM145 was predicted using AlphaFold3^25^. For the other GOLD domain–containing seven-transmembrane domain proteins, predicted structures were obtained by downloading the corresponding models from the AlphaFold2 proteome database^28, 29^. Three-dimensional images of the proteins were generated and visualized using UCSF ChimeraX^63^.

### Statistical analysis

Data analyses were performed using GraphPad Prism 9.4.1 (GraphPad Software, San Diego, CA, USA), and Excel (Microsoft, Redmond, WA, USA). Two-sided unpaired t-tests were performed to compare independent groups. For multiple-group and multiple-condition comparisons, two-way analysis of variance (ANOVA) with Dunnett’s post-hoc test was used. Statistical significance was set at p < 0.05. The number of biological replicates was indicated whenever possible. At least three independent animal or biochemical experiments were performed.

## ACKNOWLEDGEMENTS

The authors thank the Yonsei Advanced Imaging Center and Carl Zeiss Microscope. The authors also thank Medical Illustration & Design (MID), a member of the Medical Research Support Services of the Yonsei University College of Medicine, for providing excellent support with graphical illustrations.

## AUTHOR CONTRIBUTIONS

H.Y.G., C.H.K., and J.B. conceived and supervised the project. H.Y.G., C.H.K., J.B., J.W.R., and K.S.O. designed the experiments. J.W.R. performed biochemical studies, structural analysis, electrophysiological experiments, single cell RNA sequencing analysis, and mouse phenotype analysis. K.S.O. performed fluorescence microscope imaging, spatial transcriptomics analysis, and mouse phenotype analysis. J.L. and Y.C. performed fluorescence microscope imaging. S.K. conducted electron microscope imaging. J.W.H. conducted electron microscope imaging and spatial transcriptomics analysis. Y.K., H.L., and S.H.J. performed mouse studies. H.S.S. and K.Y.S. performed mouse studies and contributed to mouse phenotype analysis. J.O. and H.J. contributed to biochemical studies. J.W.R., K.S.O., C.H.K., J.B., and H.Y.G. wrote the initial manuscript. All authors reviewed and edited the manuscript.

