## Supplemental Figures 1-11 for "TMEM145 is a key component in stereociliary link structures of outer hair cells"

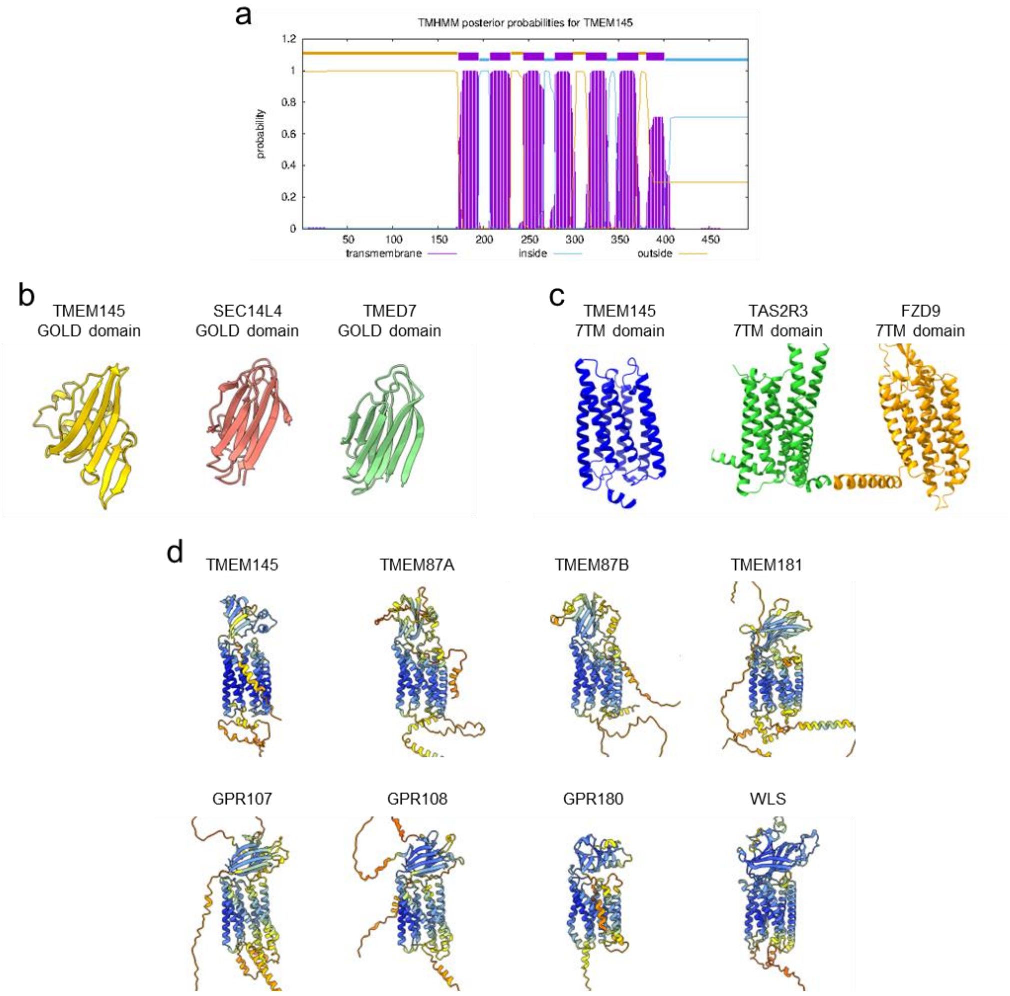


**Supplementary Fig. 1. Sequence and structure analysis of TMEM145.**

**a** Predicted transmembrane domains of TMEM145 by TMHMM 2.0

**b–d** Dali server results for GOLD domain (**b**), seven-transmembrane domain (**c**), and full-length TMEM145 (**d**).

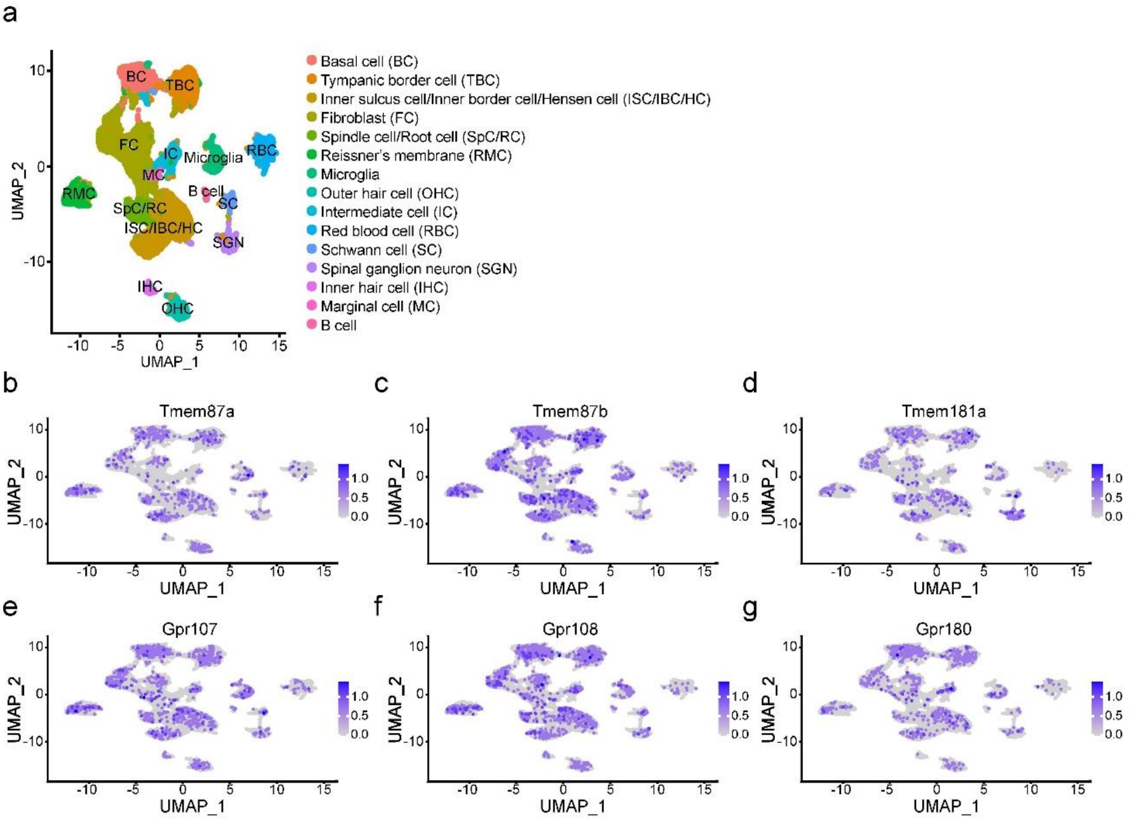


**Supplementary Fig. 2. Single cell RNA sequencing (scRNA-seq) analysis of mice cochlea at P28.**

**a** UMAP visualization of single-cell transcriptomes from the cochlea. Each point corresponds to an individual cell, color-coded by cell type. The identified cell types include Basal cell (BC), Tympanic border cell (TBC), Inner sulcus cell/Inner border cell/Hensen cell (ISC/IBC/HC), Fibroblast (FC), Spindle cell/Root cell (SpC/RC), Reissner's membrane (RMC), Marginal cell (MC), Microglia, Outer hair cell (OHC), Intermediate cell (IC), Red blood cell (RBC), Schwann cell (SC), Spiral ganglion neuron (SGN), Inner hair cell (IHC), and B cell.

**b–g** UMAP feature plots showing expression levels of *Tmem87a*, *Tmem87b*, *Tmem181a*, *Gpr107*, *Gpr108,* and *Gpr180*. The intensity of the purple color indicates relative expression (from low in gray to high in deep purple).

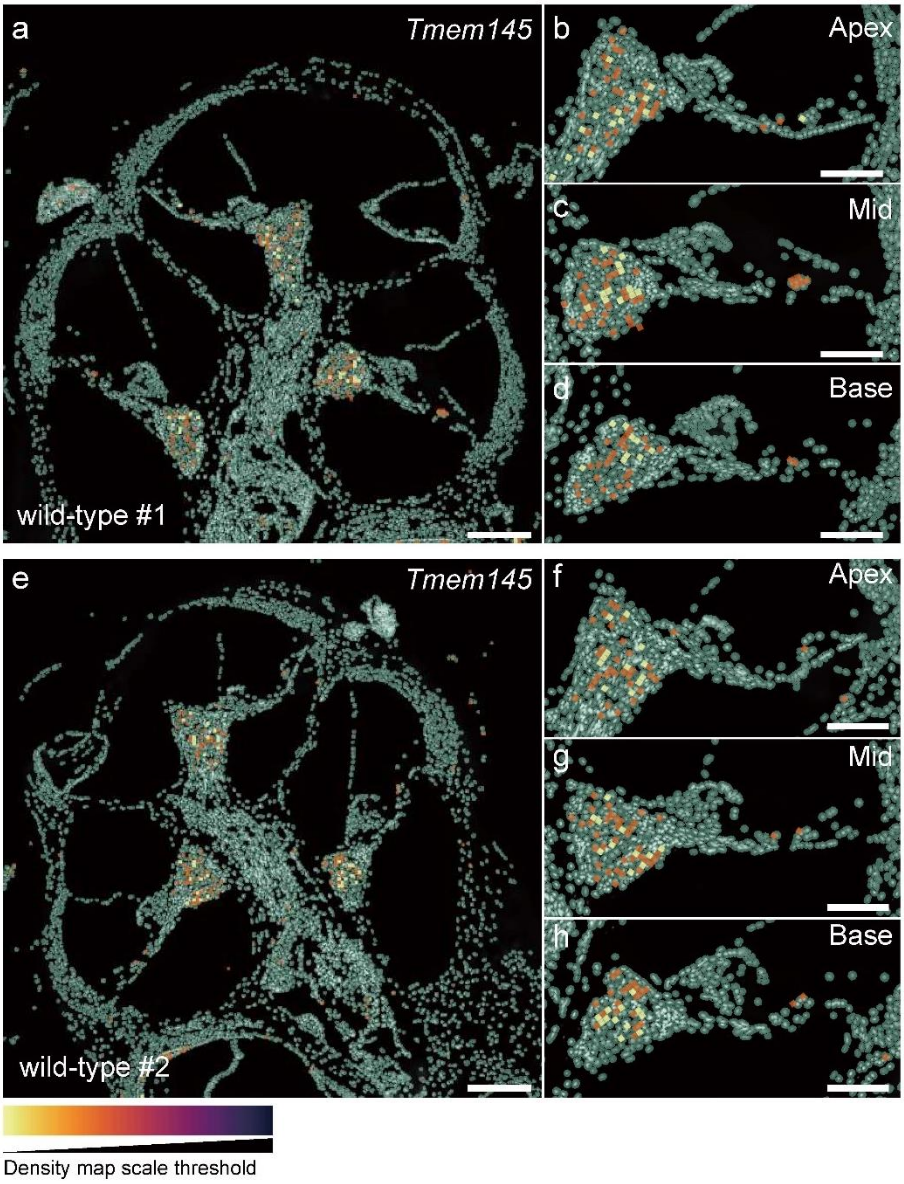


**Supplementary Fig. 3. Spatial transcriptomics analysis of *Tmem145* expression.**

**a, e** Xenium-based spatial transcriptomics analysis of the two adult mouse cochleae at P70. The color boxes indicate *Tmem145* expression.

**b–d, f–h** Spatial distribution of *Tmem145* expression in the apex, mid, and base regions of the organ of Corti.

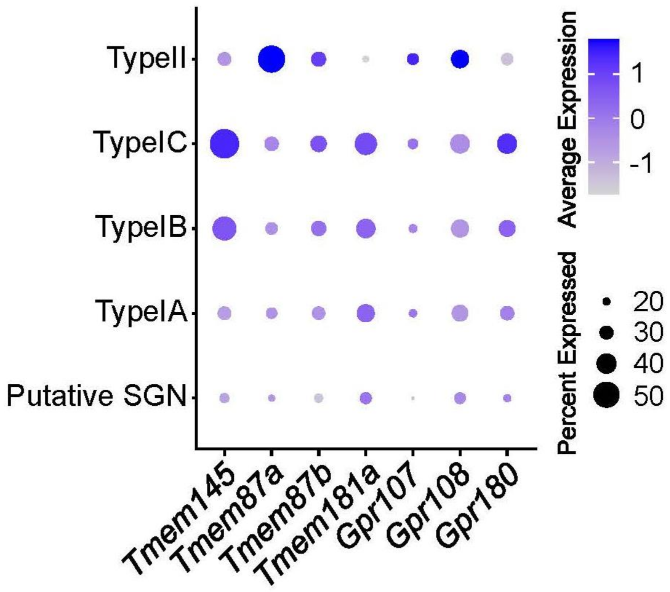


**Supplementary Fig. 4. Single cell RNA sequencing (scRNA-seq) analysis of SGN subtypes.**

Dot plot showing the expression levels of *Tmem145*, *Tmem87a*, *Tmem87b*, *Tmem181a*, *Gpr107*, *Gpr108*, and *Gpr180* across Type IA, Type IB, Type IC, Type II spiral ganglion neurons (SGNs), and putative SGN. The size of each dot reflects the percentage of cells expressing the indicated gene (see bubble size key on the right), and the color intensity denotes the average expression level (from −1 in gray to +1 in deep blue). Darker shades of blue indicate higher average expression. Each row corresponds to a different SGN subtype or putative SGN population, while each column represents one of the genes under analysis.

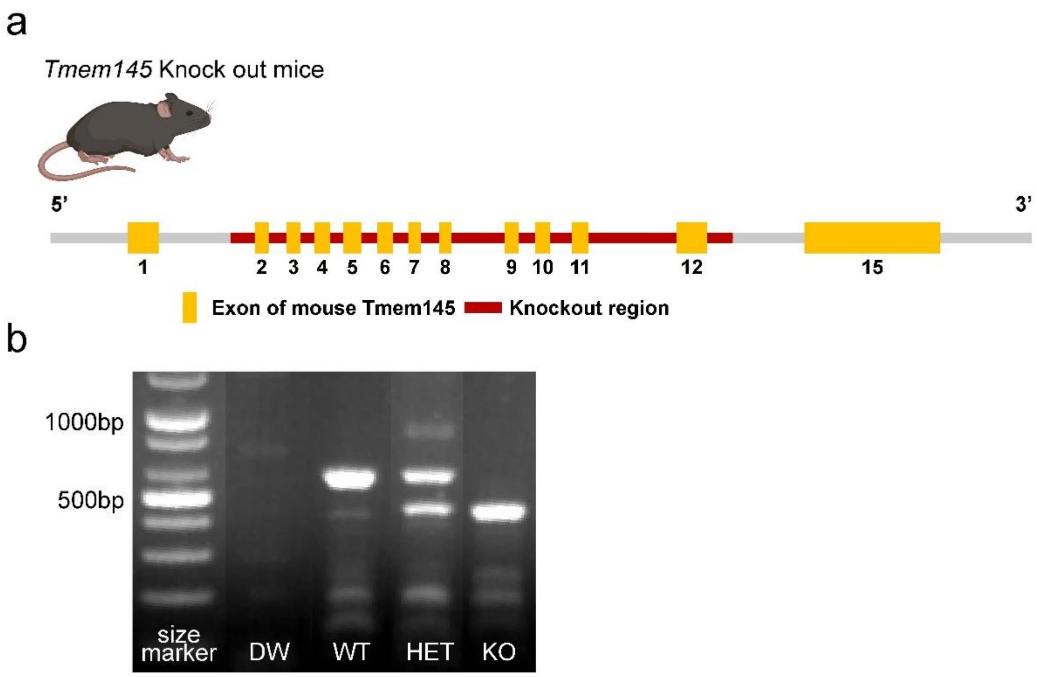


**Supplementary Fig. 5.** ***Tmem145*** **knock out mice.**

**a** Strategy for generation of *Tmem145* knock out (*Tmem145*$^{-/-}$) mice. Exons 2 to 12 of *Tmem145* in the C57BL6/N were deleted using CRISPR-Cas9.

**b** PCR genotyping result for *Tmem145*$^{-/-}$ mice. DW, distilled water, WT, wild-type (*Tmem145*$^{+/+}$), Het, Heterozygous (*Tmem145*$^{+/-}$), KO, knock out (*Tmem145*$^{-/-}$). Following primers were used: F1, 5'-GTTGCATGGTGGCTGTCGAG-3'; F2, 5'-CCTATACTCTCTGGTGAGACATGG-3'; R1, 5'-GCTGTAAATACTTCTGATTTGGTTCGG-3'

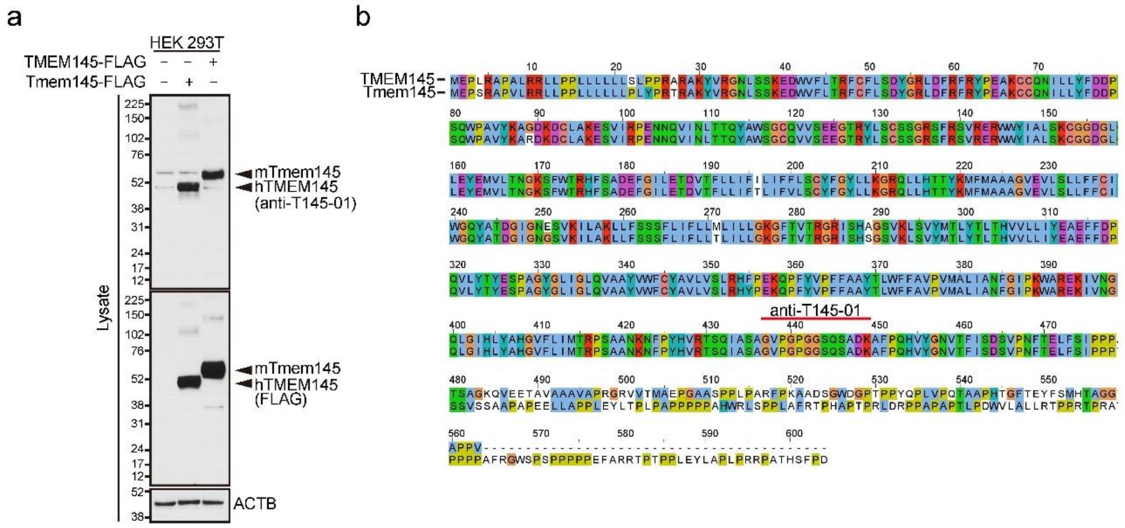


**Supplementary Fig. 6. TMEM145 antibody.**

**a** Antibody validation by Western blot from TMEM145-overexpressing HEK293T cells. The upper blot shows the result of the anti-T145-01 antibody, which was developed in this study.

**b** Amino acid sequence alignment of human TMEM145 and mouse Tmem145 proteins with the antibody binding site of anti-T145-01 indicated.

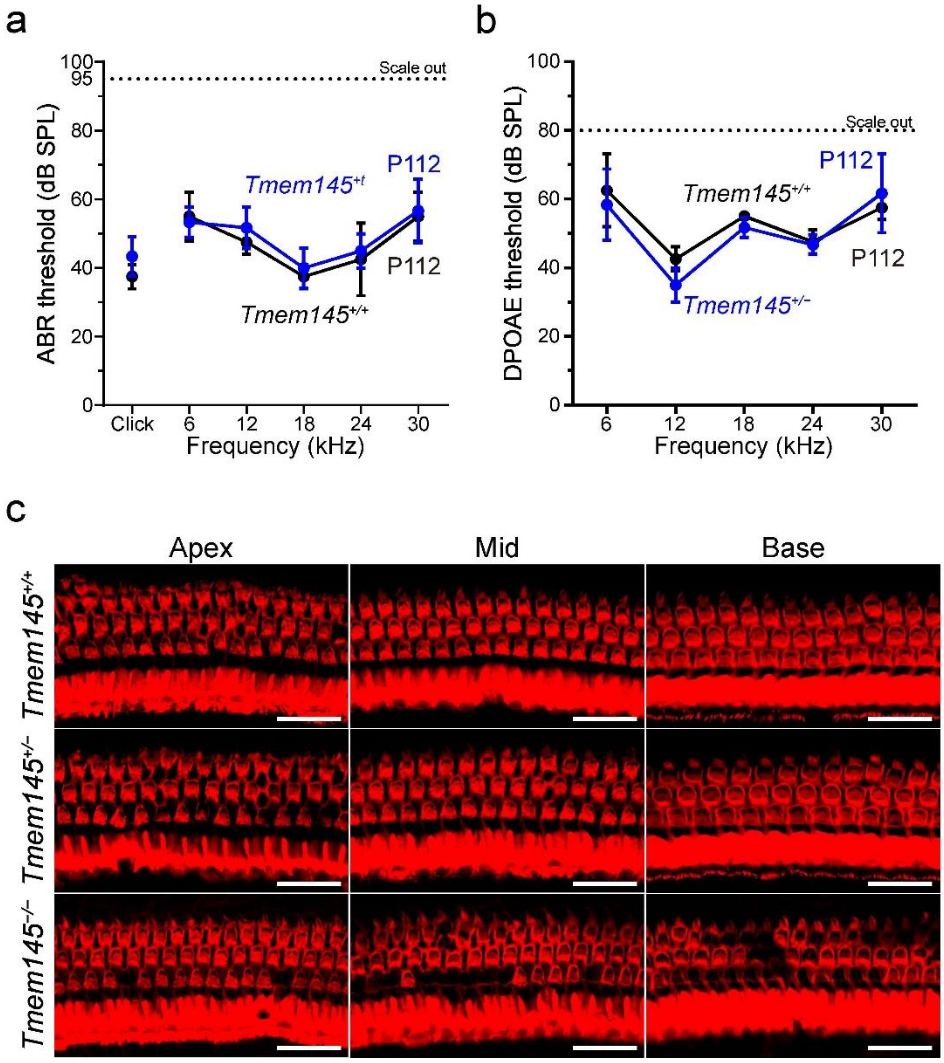


**Supplementary Fig. 7. Auditory function and hair cell survival of *Tmem145*$^{+/+}$, *Tmem145*$^{+/–}$, and *Tmem145*$^{–/–}$ mice.**

**a, b** ABR and DPOAE thresholds in *Tmem145*$^{+/+}$ and *Tmem145*$^{+/–}$ mice at P112. Compared to wild-type mice, *Tmem145*$^{+/–}$ mice exhibited no significant difference in hearing thresholds.

**c** Whole-mount immunofluorescence images of the organ of Corti with a phalloidin (red) in *Tmem145*$^{+/+}$, *Tmem145*$^{+/–}$, and *Tmem145*$^{–/–}$ mice at P112. Compared to wild-type mice, *Tmem145*$^{+/–}$ mice exhibited no significant difference in hair cell survival, but *Tmem145*$^{–/–}$ showed significant degeneration of outer hair cells (scale bars: 20 μm).

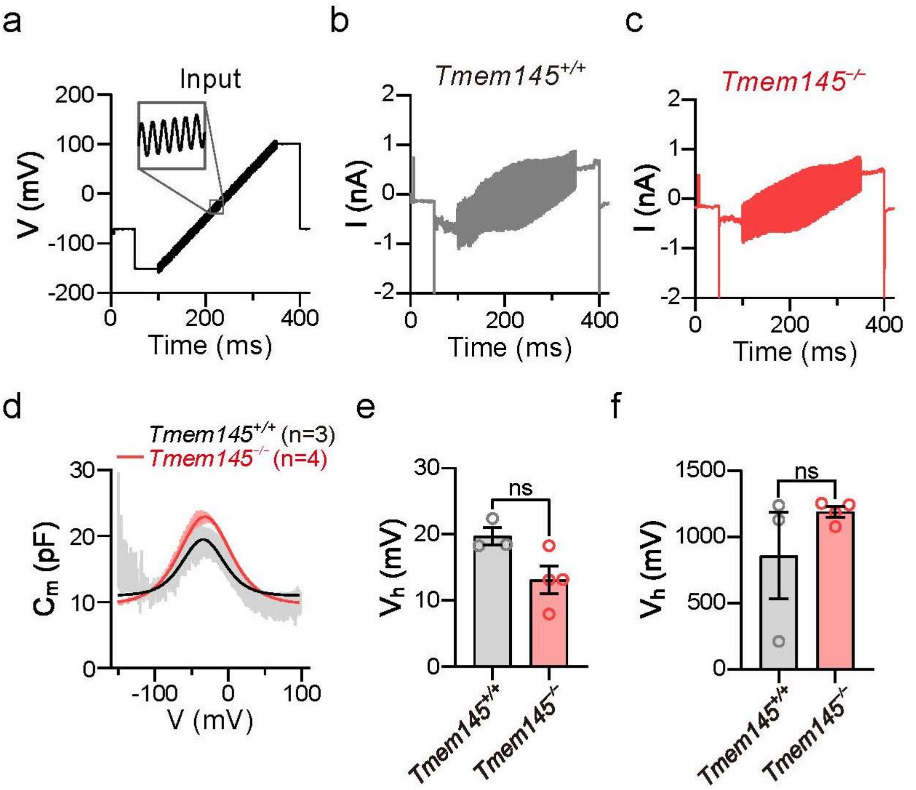


**Supplementary Fig. 8. Prestin activity in *Tmem145$^{+/+}$ and Tmem145$^{-/-}$* OHCs at P21.**

**a** Schematic of the voltage protocol used for patch-clamp recordings. The cell is held at a negative potential and then subjected to a voltage ramp (black trace), with a superimposed sinusoidal input (inset).

**b, c** Representative whole-cell current responses evoked by the protocol in (a) from *Tmem145*$^{+/+}$ (**b**, gray) and *Tmem145*$^{-/-}$ (**c**, red) OHCs.

**d** Plot of membrane capacitance ($C_m$) as a function of voltage for *Tmem145*$^{+/+}$ (gray, n = 3) and *Tmem145*$^{-/-}$ (red, n = 4) OHCs. Shaded areas indicate the standard error of the mean (SEM).

**e**, **f** Bar graphs of half-activation voltages ($V_h$) for *Tmem145*$^{+/+}$ and *Tmem145*$^{-/-}$ OHCs. Student's t-test was used to calculate the P-value with 0.05 threshold. No significant differences (ns) are observed between genotypes.

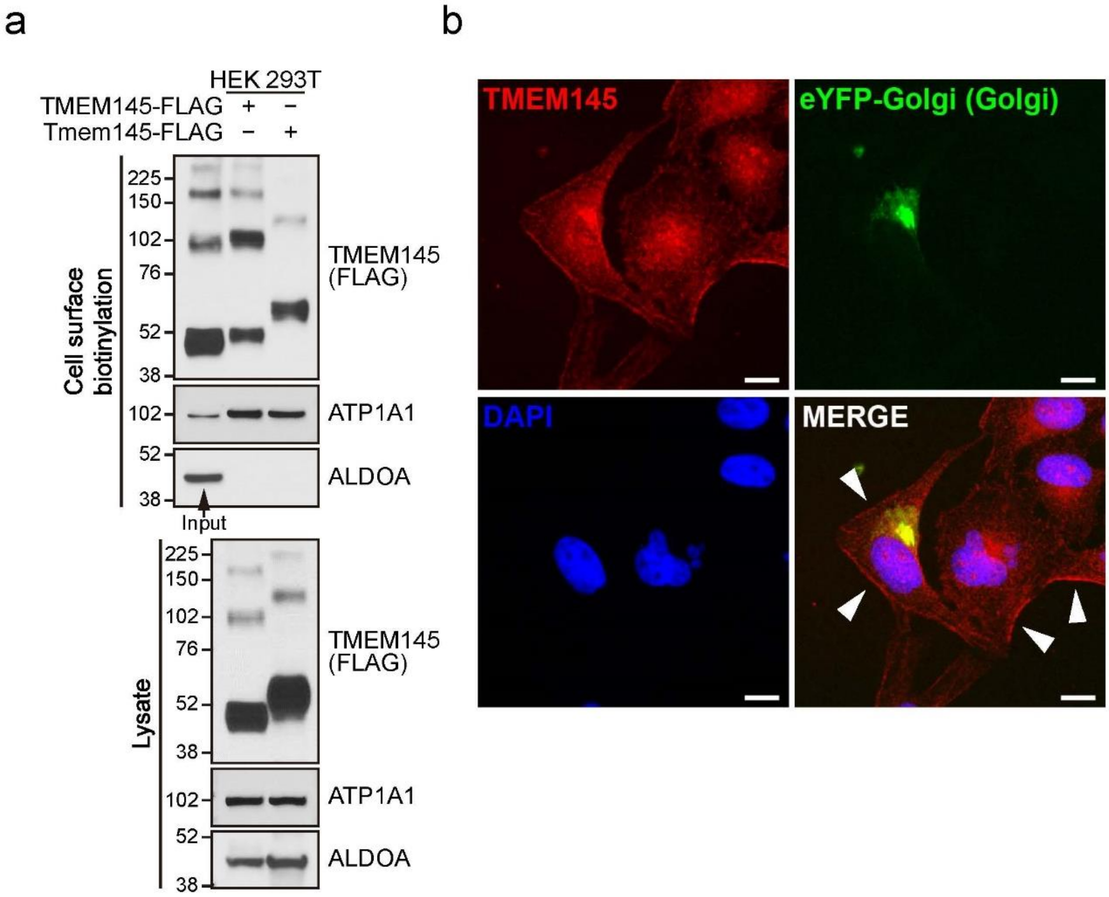


**Supplementary Fig. 9. Plasma membrane expression of TMEM145.**

**a** Cell surface biotinylation of HEK 293T cells overexpressing human TMEM145 and mouse Tmem145. Membrane proteins were labeled with biotin, isolated with avidin beads. Western blot result indicated that TMEM145 and Tmem145 were expressed on the plasma membrane. **b** Immunofluorescence of TMEM145 in HeLa cells. Arrows indicate the membrane expression of TMEM145 proteins.

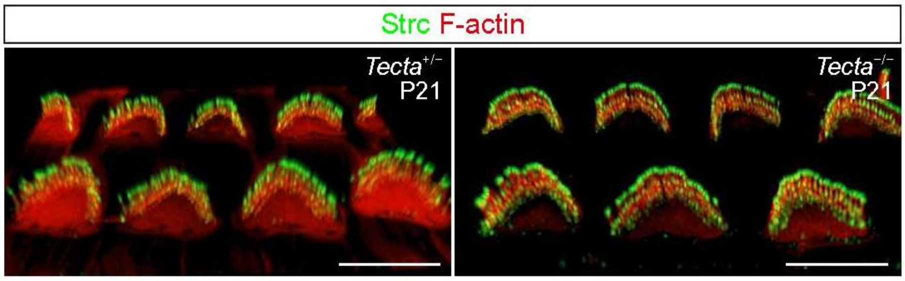


**Supplementary Fig. 10. Stereocilin localization in the stereocilia of *Tecta*$^{+/–}$ and *Tecta*$^{–/–}$ mice at P21.**

Whole-mount immunofluorescence images of the organ of Corti labeled with a STRC antibody (green) and F-actin (red) in *Tecta*$^{+/–}$ and *Tecta*$^{–/–}$ mice at P21.

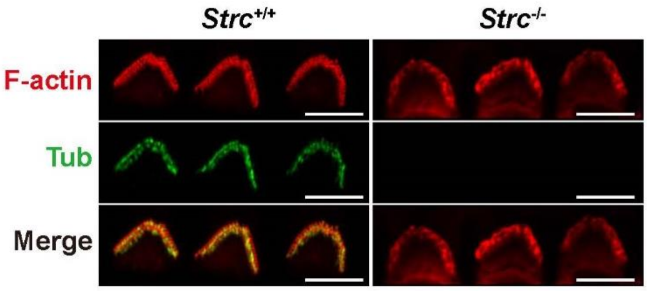


**Supplementary Fig. 11. Tubby localization in the stereocilia of *Strc*$^{+/+}$ and *Strc*$^{-/-}$ mice at P21.**

Whole-mount immunofluorescence images of the organ of Corti labeled with a TUB antibody (green) and F-actin (red) in *Strc*$^{+/+}$ and *Strc*$^{-/-}$ mice at P21.

### Supplementary Table 1. Dali server results for TMEM145 GOLD domain.

| No | Chain | Z | rmsd | align | res | %id | PDB | Description |
|---|---|---|---|---|---|---|---|---|
| 1 | e3h4-A | 22 | 0.8 | 125 | 493 | 100 | AF-Q8NBT3-F1 | TRANSMEMBRANE PROTEIN 145 |
| 2 | e75d-A | 6.9 | 2.8 | 95 | 440 | 21 | AF-Q86V85-F1 | INTEGRAL MEMBRANE PROTEIN GPR180 |
| 3 | e09y-A | 6.1 | 3.2 | 103 | 600 | 14 | AF-Q5VW38-F1 | PROTEIN GPR107 |
| 4 | e8ss-A | 6 | 3 | 101 | 543 | 15 | AF-Q9NPR9-F1 | PROTEIN GPR108 |
| 5 | fcmj-A | 4.8 | 3.8 | 82 | 1249 | 2 | AF-P29144-F1 | TRIPEPTIDYL-PEPTIDASE 2 |
| 6 | ffmr-A | 4.8 | 3 | 81 | 406 | 7 | AF-Q9UDX3-F1 | SEC14-LIKE PROTEIN 4 |
| 7 | ez5i-A | 4.8 | 2.8 | 82 | 403 | 4 | AF-O76054-F1 | SEC14-LIKE PROTEIN 2 |
| 8 | fbf9-A | 4.4 | 3.5 | 96 | 719 | 3 | AF-Q6ZSI9-F1 | CALPAIN-12 |
| 9 | fc3j-A | 4.4 | 2.6 | 68 | 224 | 6 | AF-Q9Y3B3-F1 | TRANSMEMBRANE EMP24 DOMAIN-CONTAINING PROTEIN 7 |
| 10 | e785-A | 4.3 | 2.5 | 67 | 217 | 3 | AF-Q9Y3Q3-F1 | TRANSMEMBRANE EMP24 DOMAIN-CONTAINING PROTEIN 3 |
| 11 | e9jy-A | 4.3 | 3.1 | 84 | 400 | 2 | AF-Q9UDX4-F1 | SEC14-LIKE PROTEIN 3 |
| 12 | fbpg-A | 4.3 | 3.3 | 89 | 998 | 9 | AF-P54753-F1 | EPHRIN TYPE-B RECEPTOR 3 |
| 13 | ferd-A | 4.3 | 3.5 | 86 | 640 | 3 | AF-O15484-F1 | CALPAIN-5 |
| 14 | ezb0-A | 4.3 | 3.6 | 96 | 984 | 13 | AF-P54762-F1 | EPHRIN TYPE-B RECEPTOR 1 |
| 15 | fexi-A | 4.2 | 3 | 75 | 1103 | 7 | AF-Q9H324-F1 | A DISINTEGRIN AND METALLOPROTEINASE WITH THROMBOSPONDIN MOTIFS 10 |
| 16 | e9nf-A | 4 | 3.4 | 93 | 1055 | 11 | AF-P29323-F1 | EPHRIN TYPE-B RECEPTOR 2 |
| 17 | e6oy-A | 4 | 2.8 | 72 | 240 | 6 | AF-Q8WW62-F1 | TRANSMEMBRANE EMP24 DOMAIN-CONTAINING PROTEIN 6 |
| 18 | e5vf-A | 4 | 3.3 | 77 | 827 | 8 | AF-Q9NXL6-F1 | SID1 TRANSMEMBRANE FAMILY MEMBER 1 |
| 19 | e90b-A | 3.9 | 3.1 | 81 | 813 | 1 | AF-Q9Y6W3-F1 | CALPAIN-7 |
| 20 | fd8b-A | 3.9 | 3.5 | 85 | 672 | 5 | AF-Q9HC96-F1 | CALPAIN-10 |
| 21 | e5t0-A | 3.9 | 2.4 | 64 | 227 | 6 | AF-Q13445-F1 | TRANSMEMBRANE EMP24 DOMAIN-CONTAINING PROTEIN 1 |
| 22 | e8st-A | 3.9 | 3.6 | 92 | 976 | 8 | AF-P29317-F1 | EPHRIN TYPE-A RECEPTOR 2 |
| 23 | fbmx-A | 3.9 | 3.5 | 75 | 1762 | 11 | AF-Q8N6G6-F1 | ADAMTS-LIKE PROTEIN 1 |
| 24 | e9q1-A | 3.9 | 3.7 | 92 | 983 | 10 | AF-P29320-F1 | EPHRIN TYPE-A RECEPTOR 3 |
| 25 | e0ec-A | 3.9 | 3.6 | 95 | 1037 | 8 | AF-P54756-F1 | EPHRIN TYPE-A RECEPTOR 5 |
| 26 | ffjz-A | 3.8 | 3.7 | 93 | 986 | 9 | AF-P54764-F1 | EPHRIN TYPE-A RECEPTOR 4 |
| 27 | e87v-A | 3.8 | 3.4 | 89 | 714 | 2 | AF-P07384-F1 | CALPAIN-1 CATALYTIC SUBUNIT |
| 28 | e1n8-A | 3.8 | 3.3 | 75 | 1686 | 13 | AF-Q9UKP4-F1 | A DISINTEGRIN AND METALLOPROTEINASE WITH THROMBOSPONDIN MOTIFS 7 |
| 29 | e98e-A | 3.8 | 3.5 | 77 | 1211 | 9 | AF-O95450-F1 | A DISINTEGRIN AND METALLOPROTEINASE WITH THROMBOSPONDIN MOTIFS 2 |
| 30 | ezo5-A | 3.8 | 3.5 | 94 | 1036 | 6 | AF-Q9UF33-F1 | EPHRIN TYPE-A RECEPTOR 6 |

**Supplementary Table 2. Dali server results for TMEM145 seven-transmembrane domain.**

| No | Chain | Z | rmsd | align | res | %id | PDB | Description |
|---|---|---|---|---|---|---|---|---|
| 1 | e3h4-A | 36 | 1.1 | 254 | 493 | 100 | AF-Q8NBT3-F1 | TRANSMEMBRANE PROTEIN 145 |
| 2 | e1r2-A | 22.5 | 2.8 | 234 | 555 | 12 | AF-Q96K49-F1 | TRANSMEMBRANE PROTEIN 87B |
| 3 | fgxs-A | 21.9 | 2.8 | 232 | 555 | 16 | AF-Q8NBN3-F1 | TRANSMEMBRANE PROTEIN 87A |
| 4 | e09y-A | 20.8 | 3.1 | 236 | 600 | 13 | AF-Q5VW38-F1 | PROTEIN GPR107 |
| 5 | e75d-A | 20.7 | 2.9 | 228 | 440 | 18 | AF-Q86V85-F1 | INTEGRAL MEMBRANE PROTEIN GPR180 |
| 6 | e8ss-A | 19.8 | 3 | 230 | 543 | 13 | AF-Q9NPR9-F1 | PROTEIN GPR108 |
| 7 | e9nr-A | 15.5 | 3.7 | 219 | 316 | 9 | AF-Q9NYW6-F1 | TASTE RECEPTOR TYPE 2 MEMBER 3 |
| 8 | e9ow-A | 15.4 | 3.7 | 221 | 609 | 9 | AF-Q6NV75-F1 | PROBABLE G-PROTEIN COUPLED RECEPTOR 153 |
| 9 | e8op-A | 15.4 | 3.9 | 224 | 839 | 9 | AF-Q8TE23-F1 | TASTE RECEPTOR TYPE 1 MEMBER 2 |
| 10 | fg8p-A | 15.3 | 3.9 | 226 | 303 | 9 | AF-Q9NYV9-F1 | TASTE RECEPTOR TYPE 2 MEMBER 13 |
| 11 | e6e6-A | 15.3 | 4.3 | 222 | 299 | 8 | AF-P59544-F1 | TASTE RECEPTOR TYPE 2 MEMBER 50 |
| 12 | fbmq-A | 15.2 | 3.5 | 211 | 612 | 9 | AF-Q9P2C4-F1 | TRANSMEMBRANE PROTEIN 181 |
| 13 | e22v-A | 14.9 | 4.4 | 224 | 362 | 10 | AF-P25106-F1 | ATYPICAL CHEMOKINE RECEPTOR 3 |
| 14 | e215-A | 14.9 | 3.9 | 211 | 332 | 9 | AF-P32245-F1 | MELANOCORTIN RECEPTOR 4 |
| 15 | e85s-A | 14.9 | 3.8 | 213 | 591 | 11 | AF-O00144-F1 | FRIZZLED-9 |
| 16 | e3hb-A | 14.9 | 4 | 221 | 588 | 8 | AF-Q16538-F1 | PROBABLE G-PROTEIN COUPLED RECEPTOR 162 |
| 17 | e1kl-A | 14.8 | 3.8 | 219 | 311 | 7 | AF-Q6TCH7-F1 | PROGESTIN AND ADIPOQ RECEPTOR FAMILY MEMBER 3 |
| 18 | e8cf-A | 14.6 | 3.3 | 196 | 238 | 5 | AF-Q15546-F1 | MONOCYTE TO MACROPHAGE DIFFERENTIATION FACTOR |
| 19 | fcal-A | 14.6 | 3.9 | 217 | 537 | 7 | AF-Q9ULV1-F1 | FRIZZLED-4 |
| 20 | fglo-A | 14.6 | 3.9 | 214 | 647 | 6 | AF-Q9UP38-F1 | FRIZZLED-1 |
| 21 | e2kc-A | 14.6 | 3.9 | 213 | 574 | 6 | AF-O75084-F1 | FRIZZLED-7 |
| 22 | fedl-A | 14.5 | 3.6 | 213 | 926 | 9 | AF-Q5T6X5-F1 | G-PROTEIN COUPLED RECEPTOR FAMILY C GROUP 6 MEMBER A |
| 23 | e5oc-A | 14.5 | 3.9 | 213 | 706 | 7 | AF-O60353-F1 | FRIZZLED-6 |
| 24 | fgkk-A | 14.5 | 3.9 | 215 | 565 | 7 | AF-Q14332-F1 | FRIZZLED-2 |
| 25 | fb2j-A | 14.5 | 4.1 | 213 | 342 | 10 | AF-Q9H244-F1 | P2Y PURINOCEPTOR 12 |
| 26 | e7lw-A | 14.4 | 3.8 | 209 | 814 | 8 | AF-Q8NFN8-F1 | PROBABLE G-PROTEIN COUPLED RECEPTOR 156 |
| 27 | e9m8-A | 14.4 | 4.1 | 223 | 787 | 9 | AF-Q99835-F1 | SMOOTHENED HOMOLOG |
| 28 | feno-A | 14.4 | 4.4 | 221 | 695 | 10 | AF-P23945-F1 | FOLLICLE-STIMULATING HORMONE RECEPTOR |
| 29 | e9j2-A | 14.4 | 3.9 | 219 | 319 | 10 | AF-A6NGY5-F1 | OLFACTORY RECEPTOR 51F1 |
| 30 | e6eg-A | 14.4 | 4.1 | 221 | 764 | 8 | AF-P16473-F1 | THYROTROPIN RECEPTOR |

### Supplementary Table 3. Dali server results for full-length TMEM145.

| No | Chain | Z | rmsd | align | res | %id | PDB | Description |
|---|---|---|---|---|---|---|---|---|
| 1 | e3h4-A | 41.5 | 6.9 | 452 | 493 | 95 | AF-Q8NBT3-F1 | TRANSMEMBRANE PROTEIN 145 |
| 2 | fgxs-A | 21.5 | 3.8 | 363 | 555 | 13 | AF-Q8NBN3-F1 | TRANSMEMBRANE PROTEIN 87A |
| 3 | e1r2-A | 21.5 | 6.5 | 361 | 555 | 12 | AF-Q96K49-F1 | TRANSMEMBRANE PROTEIN 87B |
| 4 | e75d-A | 20.4 | 4.1 | 355 | 440 | 20 | AF-Q86V85-F1 | INTEGRAL MEMBRANE PROTEIN GPR180 |
| 5 | e09y-A | 19.9 | 7.8 | 388 | 600 | 12 | AF-Q5VW38-F1 | PROTEIN GPR107 |
| 6 | e8ss-A | 18.5 | 7.8 | 356 | 543 | 12 | AF-Q9NPR9-F1 | PROTEIN GPR108 |
| 7 | e1kl-A | 14.8 | 4.8 | 245 | 311 | 7 | AF-Q6TCH7-F1 | PROGESTIN AND ADIPOQ RECEPTOR FAMILY MEMBER 3 |
| 8 | e9nr-A | 14.6 | 12.8 | 233 | 316 | 9 | AF-Q9NYW6-F1 | TASTE RECEPTOR TYPE 2 MEMBER 3 |
| 9 | fbmq-A | 14.5 | 9.3 | 265 | 612 | 11 | AF-Q9P2C4-F1 | TRANSMEMBRANE PROTEIN 181 |
| 10 | fg8p-A | 14.4 | 3.9 | 229 | 303 | 9 | AF-Q9NYV9-F1 | TASTE RECEPTOR TYPE 2 MEMBER 13 |
| 11 | e9ow-A | 14.4 | 16.4 | 252 | 609 | 8 | AF-Q6NV75-F1 | PROBABLE G-PROTEIN COUPLED RECEPTOR 153 |
| 12 | e5xa-A | 14.2 | 5.2 | 240 | 336 | 8 | AF-Q96P67-F1 | PROBABLE G-PROTEIN COUPLED RECEPTOR 82 |
| 13 | e6e6-A | 14.2 | 4.3 | 223 | 299 | 7 | AF-P59544-F1 | TASTE RECEPTOR TYPE 2 MEMBER 50 |
| 14 | e6f5-A | 14.1 | 8.7 | 266 | 374 | 10 | AF-O00254-F1 | PROTEINASE-ACTIVATED RECEPTOR 3 |
| 15 | fdhi-A | 14.1 | 3.8 | 237 | 386 | 9 | AF-Q86V24-F1 | ADIPONECTIN RECEPTOR PROTEIN 2 |
| 16 | fcal-A | 14.1 | 15.2 | 265 | 537 | 6 | AF-Q9ULV1-F1 | FRIZZLED-4 |
| 17 | e8op-A | 14 | 5 | 247 | 839 | 9 | AF-Q8TE23-F1 | TASTE RECEPTOR TYPE 1 MEMBER 2 |
| 18 | e8cf-A | 14 | 3.6 | 208 | 238 | 6 | AF-Q15546-F1 | MONOCYTE TO MACROPHAGE DIFFERENTIATION FACTOR |
| 19 | e7l4-A | 14 | 4.7 | 242 | 319 | 10 | AF-O14626-F1 | PROBABLE G-PROTEIN COUPLED RECEPTOR 171 |
| 20 | e85s-A | 13.9 | 6.4 | 264 | 591 | 10 | AF-O00144-F1 | FRIZZLED-9 |
| 21 | e2kc-A | 13.9 | 7.4 | 256 | 574 | 7 | AF-O75084-F1 | FRIZZLED-7 |
| 22 | e22v-A | 13.9 | 11.2 | 258 | 362 | 10 | AF-P25106-F1 | ATYPICAL CHEMOKINE RECEPTOR 3 |
| 23 | e7lw-A | 13.9 | 14.1 | 280 | 814 | 8 | AF-Q8NFN8-F1 | PROBABLE G-PROTEIN COUPLED RECEPTOR 156 |
| 24 | e3md-A | 13.8 | 3.8 | 238 | 375 | 10 | AF-Q96A54-F1 | ADIPONECTIN RECEPTOR PROTEIN 1 |
| 25 | e3hb-A | 13.7 | 16.4 | 278 | 588 | 8 | AF-Q16538-F1 | PROBABLE G-PROTEIN COUPLED RECEPTOR 162 |
| 26 | e215-A | 13.7 | 6.1 | 237 | 332 | 7 | AF-P32245-F1 | MELANOCORTIN RECEPTOR 4 |
| 27 | fb1l-A | 13.7 | 4.7 | 235 | 319 | 9 | AF-O00270-F1 | 12-(S)-HYDROXY-5,8,10,14-EICOSATETRAENOIC ACID RECEPTOR |
| 28 | fglo-A | 13.7 | 6 | 241 | 647 | 6 | AF-Q9UP38-F1 | FRIZZLED-1 |
| 29 | fdx2-A | 13.7 | 4.5 | 254 | 330 | 9 | AF-P46089-F1 | G-PROTEIN COUPLED RECEPTOR 3 |
| 30 | fbbu-A | 13.6 | 7 | 261 | 345 | 8 | AF-Q9NZD1-F1 | G-PROTEIN COUPLED RECEPTOR FAMILY C GROUP 5 MEMBER D |
